# Assessing generalizability of a dengue classifier across multiple datasets

**DOI:** 10.1101/2023.07.17.549435

**Authors:** Bingqian Lu, Yanni Li, Ciaran Evans

## Abstract

Early diagnosis of dengue fever is important for individual treatment and monitoring disease prevalence in the population. To assist diagnosis, previous studies have proposed classification models to detect dengue from symptoms and clinical measurements. However, there has been little exploration of whether existing models can be used to make predictions for new populations. We trained logistic regression models on five publicly available dengue datasets from previous studies, using three explanatory variables identified as important in prior work: age, white blood cell count, and platelet count. These five datasets were collected at different times in different locations, with a variety of disease rates and patient ages. A model was trained on each dataset, and predictive performance and model calibration was evaluated on both the original (training) dataset, and the other (test) datasets from different studies. We further compared performance with larger models and other classification methods. In-sample area under the receiver operating characteristic curve (AUC) values for the logistic regression models ranged from 0.74 to 0.89, while out-of-sample AUCs ranged from 0.55 to 0.89. Matching age ranges in training/test datasets increased AUC values and balanced the sensitivity and specificity. Adjusting the predicted probabilities to account for differences in dengue prevalence improved calibration in 20/28 training-test pairs. Results were similar when other explanatory variables were included and when other classification methods (decision trees and support vector machines) were used. The in-sample performance of the logistic regression model was consistent with previous dengue classifiers, suggesting the chosen model is a good choice in a variety of settings and has decent overall performance. However, adjustments are required to make predictions on new datasets. Practitioners can use existing dengue classifiers in new settings but should be careful with different patient ages and disease rates.

## Introduction

Dengue fever is an acute mosquito-borne infection worldwide, most common in tropical and subtropical areas, with a substantial and growing disease burden. In countries such as Vietnam, Malaysia, and Colombia, more severe dengue cases were reported in the last decades (1; 2), and the number of dengue cases doubled every 10 years from 1990 to 2013 (3). In 2019, dengue was observed in Afghanistan for the first time (4), and the World Health Organization (WHO) has also reported that dengue has been recorded in more than 100 countries worldwide and is now spreading to Europe (5; 6). Furthermore, accurate estimation of the number of cases per year is complicated by low levels of reporting, a lack of inexpensive diagnostic methods, and incompatible comparative analysis; these issues lead to underestimates of dengue cases, particularly in developing countries, and the annual number of cases may be as high as 390 million (5; 7).

### Dengue diagnosis

Dengue is characterized by symptoms such as high fever (*>* 40*°*C), muscle pain, severe headache, plasma leakage, bleeding diathesis, and skin rash (6). The standard guidelines for dengue diagnosis are given by the WHO (6), but the symptoms used for diagnosis overlap with several other diseases, such as yellow fever, Zika, and West Nile virus. It can therefore be difficult to identify dengue in the first few days of illness (8), and previous research suggests that the diagnostic guidelines need further modifications (9; 10).

Beyond general symptoms, the WHO recommends examining viral RNA and antibodies for dengue detection. Viral RNA detection offers the advantage of providing faster results, typically within 24 to 48 hours. However, viral RNA detection tools require more expensive equipment and complicated processes, which limit their use in many developing countries impacted by dengue fever (11). Furthermore, dengue antibody detection is not suitable for early diagnosis of dengue infection, particularly during primary infections. IgM antibodies are only detectable in approximately 50% of patients between day 3 and 5 from the onset of fever. The percentage of patients with detectable IgM antibodies increases to 80% by day 5 and reaches 99% by day 10. So the performance of antibody detection is the worst in the early stages of the disease (12), which is problematic because early diagnosis of dengue can be critical for timely treatment. Dengue shock syndrome (DSS), the most common life-threatening complication of dengue caused by a serious loss of blood in the circulatory system, usually onsets between the 4th and 6th day of illness, and the mortality rate can be up to 20% (8; 13). Thus, identifying dengue in the early stage is crucial for impacted countries.

According to the WHO, the gold standard of dengue diagnosis is composed of RT-PCR viral detection, IgM/IgG antibodies and NS1 detection (14). As antigens can be detected earlier, rapid antigen tests are often used together with antibody detection in early diagnosis (15), and the NS1 rapid test is generally considered very sensitive in the early stages of dengue (15). In practice, both NS1 and IgM/IgG antibodies rapid tests are widely used for diagnosis, depending on factors like the patient’s clinical presentation, the illness’s timing, and the availability of the test (16). In a study comparing the different rapid tests’ diagnostic performance, simultaneous NS1 and IgM/IgG antibodies tests gave the best performance with 92% sensitivity and 96% specificity (17). Another comparison study among RT-PCR viral detection, IgM/IgG antibodies, and NS1 detection suggests that viral RNA and NS1 antigen detection were more suitable than IgM antibodies in the early days after symptom onset. However, starting from days 4-5, IgM detection became the most effective diagnostic method. On days 6-7 and in later samples, both NS1 antigen and IgM detection outperformed viral RNA detection as diagnostic markers for dengue infection (18).

### Classification methods for diagnosis

While a combination of antigen and antibody tests performs well for detecting dengue, these rapid tests are not always available. To supplement rapid tests, previous studies have investigated the use of classifiers trained on common diagnostic measurements such as symptoms, blood work, and demographic variables. For example, Tuan *et al.* (8), Ngim *et al.* (2), and Andries *et al.* (19) recorded counts of white blood cells, platelets, and lymphocytes. Physical symptoms recorded include rashes, body temperature, bleeding, arthralgia, asthenia, cough, and headache, and demographic variables include sex and age (2; 8; 20; 21).

These symptoms, demographics, and laboratory measurements have been employed by previous studies to predict whether an ill patient actually has dengue. Tuan *et al.* (8) used logistic regression to classify patients, and found through model selection that age, white blood cell count, and platelet count were the most important predictors. Their final model has a sensitivity of 74.8% and a specificity of 76.3% for detecting dengue in children, and their results indicate that combining their classifier with rapid tests can slightly improve the performance of the rapid test. Ngim *et al.* (2) also applied logistic regression, using a penalized model to decrease small-sample bias and improve estimation. Variables were selected using stepwise selection, similar to Tuan *et al.* (8), and the final model had an area under the receiver operating characteristic (ROC) curve (AUC) of 0.82, a sensitivity of 78.4%, and a specificity of 74.6%.

Beyond logistic regression, a variety of other models have been proposed, with performance similar to Tuan *et al.* (8) and Ngim *et al.* (2). For example, Tanner *et al.* (22) employed a C4.5 decision tree classifier, using predictors such as platelet count, white blood cell count, lymphocyte count, body temperature, hematocrit count, and neutrophil count, and reported a specificity of 80.2% and a sensitivity of 78.2%. Park et al. (23) used structural equation models (SEMs), with variables also including white blood cell count, platelet cells count, age, lymphocyte counts, and hematocrit; their final model had an AUC of 0.84 at fever day-3 for dengue prediction.

### Limitations of previous studies

In a recent systematic review, Neto *et al.* (24) found limited comparisons of existing methods for dengue classification in the literature, and a dearth of available data from previous studies that would allow researchers to reproduce and compare prior results. Comparisons of previous work are further complicated by the fact that each dataset contains different explanatory variables, so a model from one study may not be usable with new data if one or more variables in the model is not recorded. Additionally, previous studies were conducted in a wide variety of locations and at different times, and it is unclear whether models built in one location and time can generalize to a new setting. To the best of our knowledge, no previous work has assessed the generalizability of dengue classification models.

### Contributions

The purpose of this paper is to evaluate whether a dengue classifier trained on one dataset can make useful predictions on a new dataset. Our work focuses on logistic regression models, which have been widely used in previous work, can be easy for practitioners to interpret (8), and which have also demonstrated their competitive performance compared to other classifiers. We have identified five publicly available datasets from previous dengue studies, which all contain commonly used explanatory variables like age, white blood cell count, and platelet count. To assess generalizability, we conduct an extensive comparison of in-sample and out-of-sample performance for models built on each of the five datasets. We further explore possible reasons a classifier may fail to generalize, and investigate adjustments to make predictions more generalizable. Finally, we consider other possible classification methods (CART classification trees and support vector machines (SVMs)), and other potential explanatory variables available in subsets of the five datasets studied here.

If dengue classifiers generalize to new data, researchers could adapt existing models for new locations, saving the substantial time and effort required to collect new data and fit a model. If classifiers do not generalize, however, then existing models have limited utility beyond the specific data used to train them.

## Methods

### Datasets

Data was used from five different papers on predicting dengue, which made their datasets publicly available. Each dataset was collected in a different location and contains varying numbers of patients and features. All raw data, and the source code for all the analysis in this manuscript, can be found in our GitHub repository

#### Dataset 1

Tuan *et al.* (8) collected data on 5726 patients, aged 1–15, who presented with possible dengue fever at seven hospitals in southern Vietnam between 2010 and 2012, with 340 patients from the smallest site and 1589 from the largest site. Of these patients, 1698 (29.7%) were diagnosed with dengue using a combination of RT-PCR, IgM serology, and NS1 detection by ELISA. Patients were eligible for the study if their fever and symptom history was less than 72 hours. Blood samples, personal information, and symptoms were collected from each patient at the time of enrolment. A total of 35 variables were recorded, including white blood cell count, platelet count, temperature, height, and weight, and symptoms such as vomiting, rashes, skin bleeding, and mucosal bleeding. We omitted two patients with missing values for white blood cell count and platelet count.

#### Dataset 2

Gasem *et al.* (25) collected data on 1486 fever patients (body temperature at least 38*°*C) aged 1–98 from eight different sites in Indonesia between July 2013 and June 2016, with 65 patients from the smallest site and 267 from the largest site. Of these patients, 467 (31.4%) were diagnosed with dengue. A total of 51 variables were recorded on the symptoms, laboratory results, and final diagnosis for each subject, and the site at which each subject was recorded. Diagnoses included Streptococcus, dengue, Chikungunya, influenza, and RSV, among others. To focus on identifying dengue fever, we grouped all non-dengue diagnoses together as one category for analysis. We omitted one patient with a missing value for platelet count.

#### Dataset 3

Saito *et al.* (21) collected data on 1573 patients aged 1–85, with a fever lasting at most 21 days and suspected of bloodstream infection, in the Philippines from 2015 to 2019. Of these patients, 256 (16.3%) were diagnosed with dengue. A total of 284 variables were recorded on the symptoms, laboratory results, medical history, and the diagnosis from laboratory tests for each subject. We omitted 21 patients with missing values for platelet count or white blood cell count, leaving 1552 rows in the data.

#### Dataset 4

Park *et al.* (23) collected data on 257 children aged 6 months to 15 years from Thailand, with fever less than 72 hours. Of these patients, 156 (60.7%) were diagnosed with dengue; of the 156 dengue patients, 51 were diagnosed with dengue hemmorhagic fever, and nine were diagnosed with dengue shock syndrome. A total of 21 variables were recorded, including the patient’s age, sex, final diagnosis, and laboratory results. Each of the laboratory results was recorded twice at different time points, one of which was four days before patients had a fever (fever day-3), and the other was two days before patients had a fever (fever day-1). In the dataset, age is recorded as a two-year range (e.g., 12–13); we converted the range to a single numerical value by taking the midpoint of the range (e.g., 12.5).

#### Dataset 5

Ngim *et al.* (2) collected data on 368 adult patients with possible dengue infection in Malaysia in 2018. Of these patients, 167 (45.4%) were ultimately diagnosed with dengue using a laboratory test. A total of 82 variables were recorded, including dengue status, white blood cell count, platelet count, age, and symptoms such as fatigue, headache, and chills.

### Choice of explanatory variables

There are many possible explanatory variables available in each dataset, and most of the available variables do not appear in all five datasets. However, previous research suggests that a small number of features can be used for dengue classification, so to assess generalizability we restricted our attention to some of the most common and widely-used features available across studies.

Using variable selection, Tuan *et al.* simplified a full model containing 17 variables with several interaction terms to a simpler, final model with only three explanatory variables: patient age, white blood cell count, and platelet count (8). Their final model with these three variables had a senstivity of 74.8%, a specificity of 76.3%, and an AUC of 0.829. Similarly, Cavailler *et al.* (26) created a logistic regression model with five variables: positive tourniquet test, absence of upper respiratory infection symptoms, platelet count, white blood cell count, and liver transaminases (sensitivity: 75.7%, specificity: 65.7%, AUC: 0.71). Tuan *et al.*’s work is also similar to the model created by Ho *et al.* (27), who created a logistic regression model with patient age, temperature, white blood cell count, and platelet count, and reported an AUC of 0.846. Ho *et al.* also compared the performance of their small logistic regression model to a larger model with 18 explanatory variables, and saw a similar AUC of 0.841. Ngim *et al.* (2) also chose a relatively small logistic regression model with seven variables, including white blood cell count and platelet count.

To assess generalizability of dengue classification, we focused on logistic regression with three explanatory variables which are available in all five of the datasets we studied: age, white blood cell count, and platelet count. These explanatory variables were also identified as important dengue predictors by Park *et al.* (23) and Potts *et al.* (28).

### Logistic regression

We focused on a logistic regression model to predict dengue status because logistic regression is familiar to many researchers, is relatively easy to interpret, works well with a small number of explanatory variables, and has been widely used in prior studies (2; 8; 26; 27).

Furthermore, logistic regression has competitive performance with other classification methods. For example, Tuan *et al.* compared logistic regression with classification and regression trees (CART) and random forests, and chose logistic regression for their final model (8). Ho *et al.* reported similar results when comparing logistic regression with decision trees and deep neural networks; the respective AUCs of the three methods were 0.797, 0.794, and 0.810 (27). The AUCs for structural equation models (SEMs) reported by Park *et al.* were similar, ranging from 0.73 to 0.94 depending on the time of illness (23).

Logistic regression was fit using R version 4.2.1 (29). All code for model fitting and analysis can be found in our GitHub repository (https://github.com/ciaran-evans/dengue-generalizability).

### Assessing in-sample performance

Before assessing generalizability of dengue predictions to different datasets, we summarized the in-sample performance of the logistic regression model (with age, white blood cell count, and platelet count as explanatory variables) on each of the five datasets. Summarizing in-sample performance allows us to better understand the expected performance of the model, which helps develop a baseline for comparison across datasets.

In-sample performance was assessed by fitting a logistic regression model on each dataset, and evaluating the model’s performance on the same dataset. Cross-validation was used to evaluate performance on new observations from each dataset. If data were collected from multiple sites, cross-validation was performed by holding out each site in turn, as in Tuan *et al.* (8). Otherwise, 10-fold cross-validation was used. Predictive performance was measured using sensitivity, specificity, positive predictive value (PPV), negative predictive value (NPV), and area under the ROC curve (AUC); these performance metrics were calculated using the predicted probabilities from cross-validation for each observation.

Sensitivity, specificity, PPV, and NPV all require that the predicted probabilities produced by the logistic regression model are converted into binary predictions, which requires a choice of threshold (AUC, on the other hand, considers *all* possible thresholds for binarization). If a particular threshold was chosen in the original study from which we obtained the data (e.g., Tuan *et al.* (8) used a threshold of 0.33), we used that threshold for consistency. If no threshold was chosen, we selected the threshold which maximized the geometric mean of sensitivity and specificity (30; 31).

### Assessing model generalizability

A model *generalizes* to a new dataset or population when it can be used to make useful predictions on new observations. In the context of many classification methods, including logistic regression, predictions on new data can be presented as a predicted *probability* (in this case, the estimated probability that the individual has dengue). For these predicted probabilities to be useful, we therefore require (a) that the predictions can distinguish dengue patients from non-dengue patients, and (b) that the estimated probability be approximately *calibrated*, i.e. close to the true probability of dengue. Here we assess both aspects of generalizable predictions, and also explore adjustments to improve calibration of the predicted probabilities.

To assess generalizability of the logistic regression model with age, white blood cell count, and platelet count as explanatory variables, we considered all possible pairs of the available datasets, with one of the datasets as the training data and one of the remaining datasets as the test data. The logistic regression model was fit on the training set, and applied to the test set to calculate a predicted probability of dengue for each patient in the test set. We then calculated AUC, sensitivity, specificity, positive predictive value, and negative predictive value for the test set using these predicted probabilities. For metrics which required a threshold for binary classification, we considered both the thresholds for the training and test data from our assessment of in-sample performance.

In practice, we usually only have the training data to train and assess the model’s performance, which means the model would make predictions and calculate the metrics using the threshold from the training data when we apply it to new data. Given information on both the training and test datasets, we examined thresholds from both the training and test datasets when calculating the sensitivity and specificity. Exploring both thresholds provides insight into whether the choice of threshold is generalizable.

#### Restricting age ranges

It is well known that statistical methods perform poorly when asked to extrapolate to new data beyond the range of the training sample. As age is an explanatory variable in our logistic regression model, and the range of patient ages differs substantially between datasets, we re-assessed model generalizability after restricting the age range in test sets.

When the training set was Dataset 1 or Dataset 4 (age *<* 16 years), and the test set was Dataset 2 or Dataset 3, we restricted the age range of the test set to also be below 16 years. When the training set was Dataset 5 (age *>* 16 years), and the test set was Dataset 2 or Dataset 3, we restricted the age range of the test set to also be above 16 ears. We re-calculated all performance metrics for predictions on the age-restricted test sets, and compared the results to predictions on the unrestricted test sets. When re-calculating the performance metric, we only use the threshold from the training data to calculate metrics that require a threshold for binary classification to evaluate whether restricting the age range would be an efficient way to improve the generalizability of the model in practice.

#### Assessing calibration

To make a binary decision about whether or not a patient has dengue, it is important that the predicted probabilities of dengue should be close to the true probabilities in the test data, in which case the model predictions are said to be *calibrated*. We assessed calibration through calibration plots, in which the predicted probabilities for the test data are binned into deciles, and the true rate of dengue cases in each bin is plotted against the bin midpoint. When predictions are calibrated, the points should be close to the diagonal line with intercept 0 and slope 1; deviations from this diagonal indicate mis-calibrated predictions. Mis-calibrated predictions do not necessarily impact AUC (because the AUC is calculated over all possible thresholds, and any monotonic transformation of the predictions will result in the same AUC), but will lead to problems when applying a threshold to binarize predicted probabilities.

#### Calibrating classifier predictions

If model predictions fail to generalize, it is because the distribution of data has changed between training and test sets. It is generally impossible to correct the predictions and improve generalizability without making assumptions about the specific nature of this distributional change. Under certain assumptions, however, it is possible that the predicted probabilities will distinguish positive and negative cases, and can be calibrated to produce good estimates of the probability of dengue in new patients.

The two common assumptions made about distributional change are *covariate shift* and *label shift*. Covariate shift (32; 33) posits a change in the distribution of the explanatory variables, but no change in the relationship between the explanatory variables and the response. If covariate shift holds, then classifier predictions should generalize without correction, because the training and test sets have the same relationship between predictors and response. Conversely, label shift (34; 35; 36) assumes that the distribution of the response (i.e., dengue prevalence) changes between datasets, but the distributions of age, white blood cell count, and platelet count remain the same conditional on dengue status. Label shift has received substantial attention in the classification and machine learning literature, and previous work has suggested that label shift may be an appropriate assumption for modeling diseases, in which the disease rate can change but the symptoms remain the same (36). Label shift may also be suitable for the datasets considered here, as the proportion of study patients who were diagnosed with dengue ranges from 16% in Dataset 3 to 61% in Dataset 4.

Under label shift, predictions on test data can be calibrated if the prevalence of dengue in the test data is known, without having to re-fit the model on the test data. Let *p̂*_*i*_ denote the predicted probability of dengue for the *i*th patient in the test set, using a model fit on a different training set. Let π_train_ denote the marginal prevalence of dengue in the training set, and π_test_ the marginal prevalence of dengue in the test set. If π_test_ is known, then the calibrated predictions *p̂*_*i*__calibrated_ on the test set can be calculated by applying Bayes’ rule:

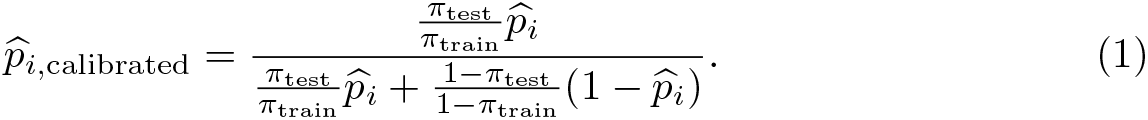

After restricting age ranges in the test set as described above, we applied Eq. (1) and assessed performance of the label shift-adjusted predictions with calibration plots. We also compared predictive performance by calculating deviance for the test data (−2×the binomial log-likelihood) using the uncorrected predictions and the label shift-adjusted predictions.

If the rate of dengue π_test_ in an area is unknown, it may be possible to estimate this rate by leveraging the label shift assumption. Lipton *et al.* (36) proposed a discretization method for estimating test prevalence under label shift, which involves inverting an estimated confusion matrix. An alternative is a fixed point method proposed by Saerens *et al.* (34), selecting the estimate of the test prevalence which most closely matches the mean of the corrected predictions from Eq. (1). We assessed performance of both methods, and compare to the simple estimate of test prevalence which averages the uncorrected predicted probabilities *p̂*_*i*_ for the test observations.

### Exploring other explanatory variables and classification methods

To assess whether predictive performance and generalizability could be improved with other explanatory variables, we identified three alternative sets of explanatory variables which were shared by at least three of the five datasets (see Table 1). For each alternative set of explanatory variables, predictive performance on all possible pairs of the available datasets was assessed, with one of the datasets as the training set and one of the remaining datasets as the test set. Performance was measured on the test set using AUC, sensitivity, specificity, positive predictive value, and negative predictive value. When the training and test sets were the same, cross-validation was used as described previously. For each training set, we compared a logistic regression model built on all variables in the alternative set; the original logistic regression model with age, white blood cell count, and platelet count; and the “best” logistic regression model selected through best subset selection with Akaike’s information criterion (AIC) (37). The `bestglm` R package was used for best subset selection with logistic regression (38).

**Table 1.** Definitions of the alternative sets of explanatory variables considered to assess in-sample performance and predictive generalizability.

| Variable subset | Explanatory variables | Datasets containing the subset |
| --- | --- | --- |
| Original | Age, White blood cell count, Platelet count | Dataset 1, Dataset 2, Dataset 3, Dataset 4 Day -3, Dataset 4 Day -1 |
| Alternative subset 1 | Sex, Age, White blood cell count, Platelet count, Lymphocytes, Neutrophils, Temperature, Hematocrit, Abdominal pain, Vomiting, Weight | Dataset 1, Dataset 3, Dataset 5 |
| Alternative subset 2 | Sex, Age, White blood cell count, Platelet count, Lymphocytes, Temperature, Blood pressure, Hemoglobin | Dataset 2, Dataset 3, Dataset 5 |
| Alternative subset 3 | Sex, Age, White blood cell count, Platelet count, Lymphocytes, Hematocrit, AST, ALT | Dataset 1, Dataset 3, Dataset 4 Day -3, Dataset 4 Day -1 |

#### Model comparison

In addition to logistic regression models, we assessed the performance of CART classification trees and support vector machines (SVMs) with a radial basis kernel (37) on all four sets of explanatory variables shown in Table 1. Performance was compared using the AUC for each pair of training and test sets. Decision trees and SVMs were fit using the `rpart` and `e1071` packages, respectively, in R (39; 40).

## Results

### Feature distributions in each dataset

The distribution of each explanatory variable in each dataset is shown in Fig. 1, which shows that these features share certain similarities in their distributions as differences. Full summary statistics for each variable are provided in Table S1. The white blood cell count and platelet count in each dataset were scaled to the same measuring unit, ×10^3^*/*mm^3^, for comparison. Despite differences in the range of age, the age distribution was unimodal and right-skewed in almost all datasets, except for Dataset 4, where the distribution of age was unimodal and roughly symmetric. White blood cell count had a right-skewed distribution that centered around 8×10^3^*/*mm^3^ among all datasets. The distribution of platelet count was also right-skewed, except in Dataset 1 and Dataset 5, in which the distributions of platelet count were unimodal and roughly symmetric. The center of the platelet count was around 200×10^3^*/*mm^3^ among the five datasets. Lastly, for Dataset 4, which collected the data of white blood cell count and platelet count from both two days and four days before the onset of fever, the distributions of white blood cell count and platelet count collected four days before the fever were more similar to the distributions we observed in other datasets. The mean of platelet count and white blood cell count from data collected two days before the fever are the lowest among the five datasets (supplementary table).

**Fig 1.**
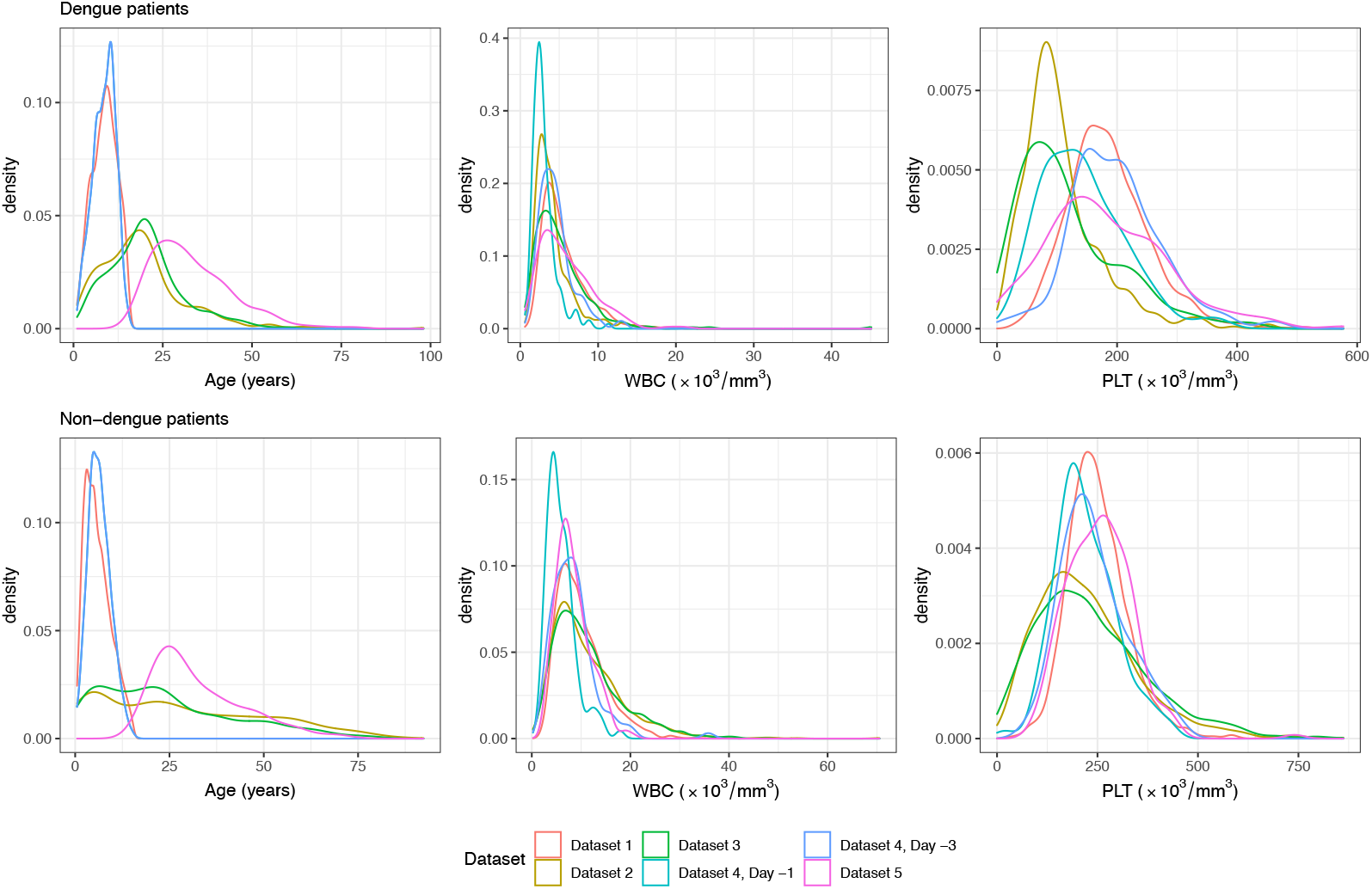
Distributions of explanatory variables across datasets. The distribution of age, white blood cell count (WBC), and platelet count (PLT) for patients in each dataset, shown separately for dengue and non-dengue patients.

### In-sample performance

The in-sample performance of each dataset is summarized in Table 2. The model on Dataset 5 had a sensitivity of 72%, a specificity of 64%, and an AUC of 0.74 (95% CI: (0.69, 0.79)), which is the lowest value among the five datasets and slightly lower than the AUC (0.82) reported by Ngim *et al.* (2). All other datasets had sensitivities and specificities around 75%, and AUCs above 0.82. The AUCs for Dataset 1 (0.83, 95% CI: (0.81, 0.84)) and Dataset 4, Day-3 (0.82, 95% CI: 0.77, 0.88)) were close to the AUCs reported by (8) and (23) (0.83 and 0.84, respectively), while the AUC for Dataset 4, Day-1 (0.86, 95% CI: (0.81, 0.90)) was slightly lower than the value of 0.93 reported by Park *et al.* (23). AUC values were not reported by Gasem *et al.* (25) and Saito *et al.* (21) for Datasets 2 and 3; we found respective AUC values of 0.89 (95% CI: (0.87, 0.91)) and 0.85 (95% CI: (0.83, 0.87)), demonstrating relatively good predictive performance.

**Table 2.** In-sample performance results for each dataset.

| Dataset | Threshold | Sensitivity | Specificity | PPV | NPV | AUC |
| --- | --- | --- | --- | --- | --- | --- |
| Dataset 1 | 0.33 | 0.74 | 0.76 | 0.56 | 0.87 | 0.83 |
| Dataset 2 | 0.45 | 0.79 | 0.85 | 0.70 | 0.90 | 0.89 |
| Dataset 3 | 0.21 | 0.78 | 0.78 | 0.41 | 0.95 | 0.85 |
| Dataset 4, Day -3 | 0.59 | 0.82 | 0.70 | 0.81 | 0.72 | 0.82 |
| Dataset 4, Day -1 | 0.64 | 0.82 | 0.77 | 0.85 | 0.74 | 0.86 |
| Dataset 5 | 0.43 | 0.72 | 0.64 | 0.62 | 0.74 | 0.74 |

### Generalizability

Model performance for each training/test pair is shown in Fig. 2, and summarized in Table S2. As expected, AUC values tended to be slightly lower than in-sample performance when using different training/test sets. However, the differences were mostly small, which supports model generalizability. The exception was training on Datasets 1 and 4, and testing on Datasets 2 and 3; these combinations had substantially lower AUCs on the test data, likely because Datasets 1 and 4 included only pediatric patients, whereas the other datasets included adult patients. The lowest AUC value, 0.547, was observed when the model was trained on Dataset 4 fever day-3 and tested on Dataset 2. However, when we tested the model on Datasets 1 and 4 using Dataset 2 and Dataset 3 as the training data, all AUC values were greater than 0.75. Datasets 2 and 3 include both pediatric and adult patients, and so extrapolation is not required to predict on Datasets 1 and 4. However, models trained on Dataset 5 (age *>* 16 years) still predicted well on Datasets 1 and 4. This may suggest that datasets with a wider age range can extrapolate to younger patients who are not in the age group more accurately.

**Fig 2.**
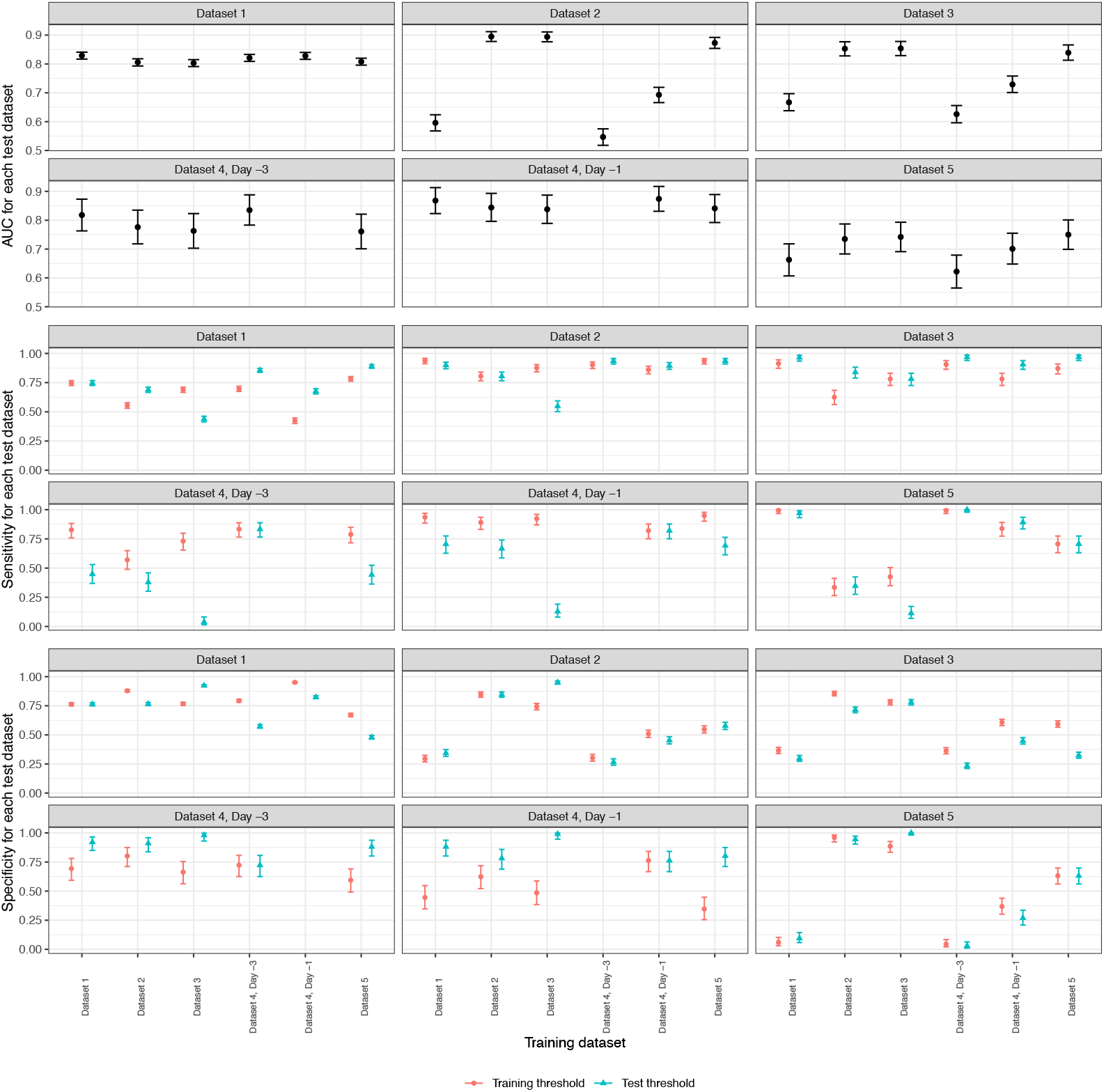
Model performance for each pair of training/test datasets. Each panel represents one test set, with predictive performance on that test set for each training set. Predictive performance is measured by AUC, sensitivity, and specificity. Sensitivity and specificity are both threshold dependent, and so we consider thresholds from both the training and test data (Table 2.)

There were also several training/test pairs with higher AUCs than the observed in-sample performance. The highest AUC value, 0.894, was observed when the model was trained on Dataset 3 and tested on Dataset 2; this AUC was higher than the in-sample performances of both Dataset 2 and Dataset 3 (Table 2). Similarly, training on Dataset 1 and testing on Dataset 4 fever day-1 also gave an AUC higher than the in-sample performances for both Dataset 1 and Dataset 4. The error bars in Fig. 2 suggest that these differences in AUC are likely due to chance.

While AUC is threshold-independent, sensitivity and specificity are not. Fig. 2 shows that using a pre-specified threshold resulted in substantial imbalance between sensitivity and specificity, suggesting that one probably needs to use an alternative way of choosing the threshold for the model to be more efficient in practice.

### Restricting age ranges

Since Dataset 2 and Dataset 3 had wider age ranges than the other datasets, we restricted their age range and re-assessed the model’s generalizability when testing on the restricted Dataset 2 and Dataset 3.

As shown in Fig. 3, restricting the age range of the test set to match the training set improved the AUC and often led to a more balanced combination of sensitivity and specificity. An exception was training on Dataset 4 fever day-1 with a threshold of 0.64, and testing on Dataset 3; the AUC improved, but the imbalance between the sensitivity and specificity was not alleviated. Since 0.64 was the largest threshold across the five datasets (and the training threshold for Dataset 3, 0.21, was the smallest), the imbalance after filtering the data may be caused by the great discrepancies between the threshold value and the proportion of dengue patients in Dataset 3.

**Fig 3.**
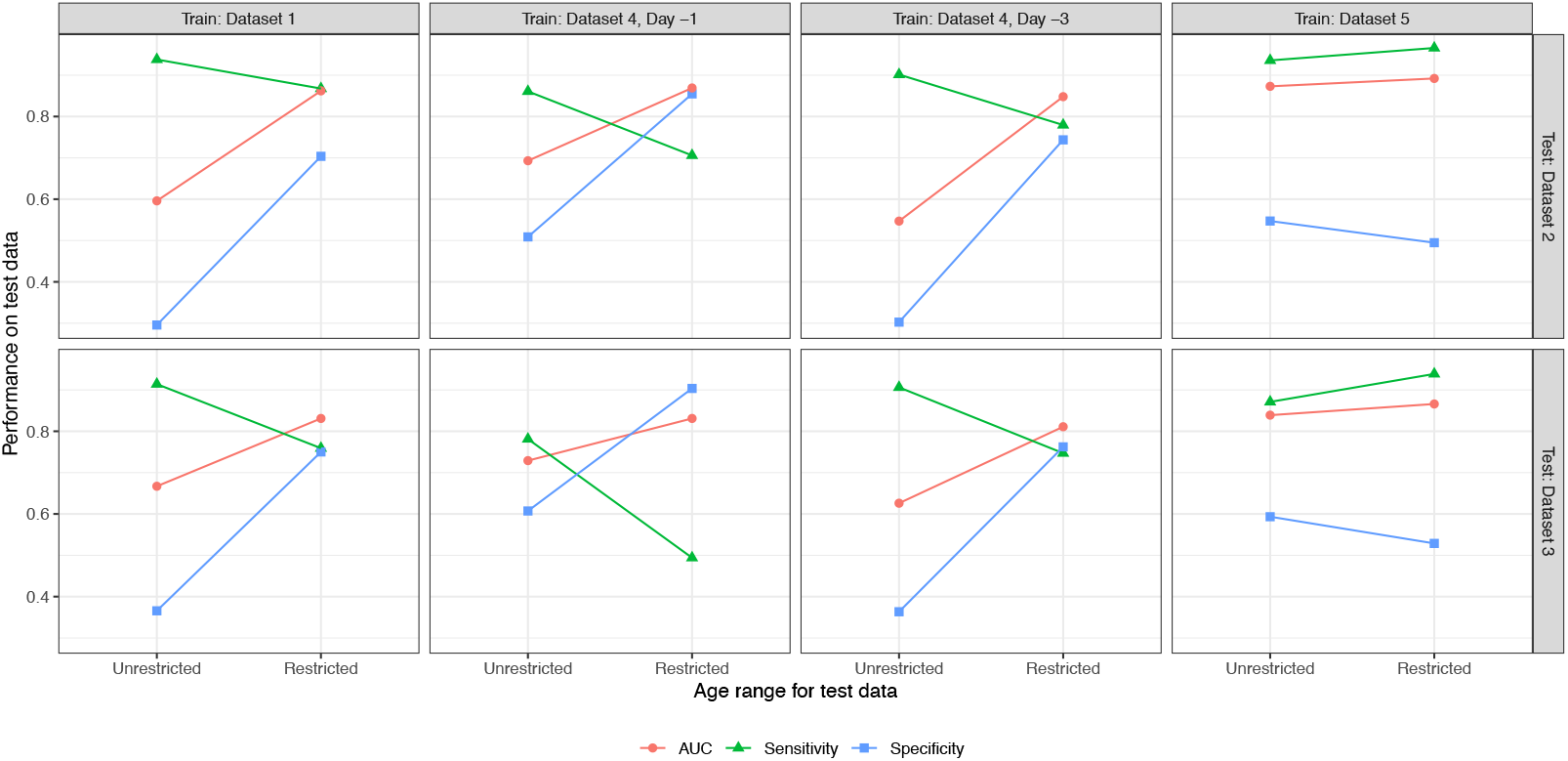
Model performance with restricted age ranges. Changes in AUC, sensitivity, and specificity are shown when restricting the age range of test data from Dataset 2 and Dataset 3. The Unrestricted performance metrics are calculated on the full test sets, while the Restricted performance metrics are calculated to match the age range in the training data (*<* 16 for Datasets 1 and 4, *>* 16 for Dataset 5). For sensitivity and specificity, the threshold for each training set (Table 2) was used.

The model trained on Dataset 5 and tested on restricted Dataset 2 and Dataset 3 showed a similar performance with improved AUC values but still imbalanced sensitivity and specificity, which may be due to the differences between the dengue proportion and the value of the chosen threshold as well.

### Calibration and label shift

Several calibration plots for different training/test pairs are shown in Fig. 4 (calibration plots for all pairs are shown in Fig. S1 and Fig. S2). The uncorrected predicted probabilities (applying the logistic regression model, fit on the training data, directly to the test data) tended to be poorly calibrated, particularly when the prevalence of dengue differed between training and test sets. Adjusting the predicted probabilities with the dengue prevalence in the test data (Eq. (1)) often improved calibration, as shown in Fig. 4. The deviance for the test data decreased in 20 out of 28 training/test pairs when using these label shift-adjusted predictions (Table S3).

**Fig 4.**
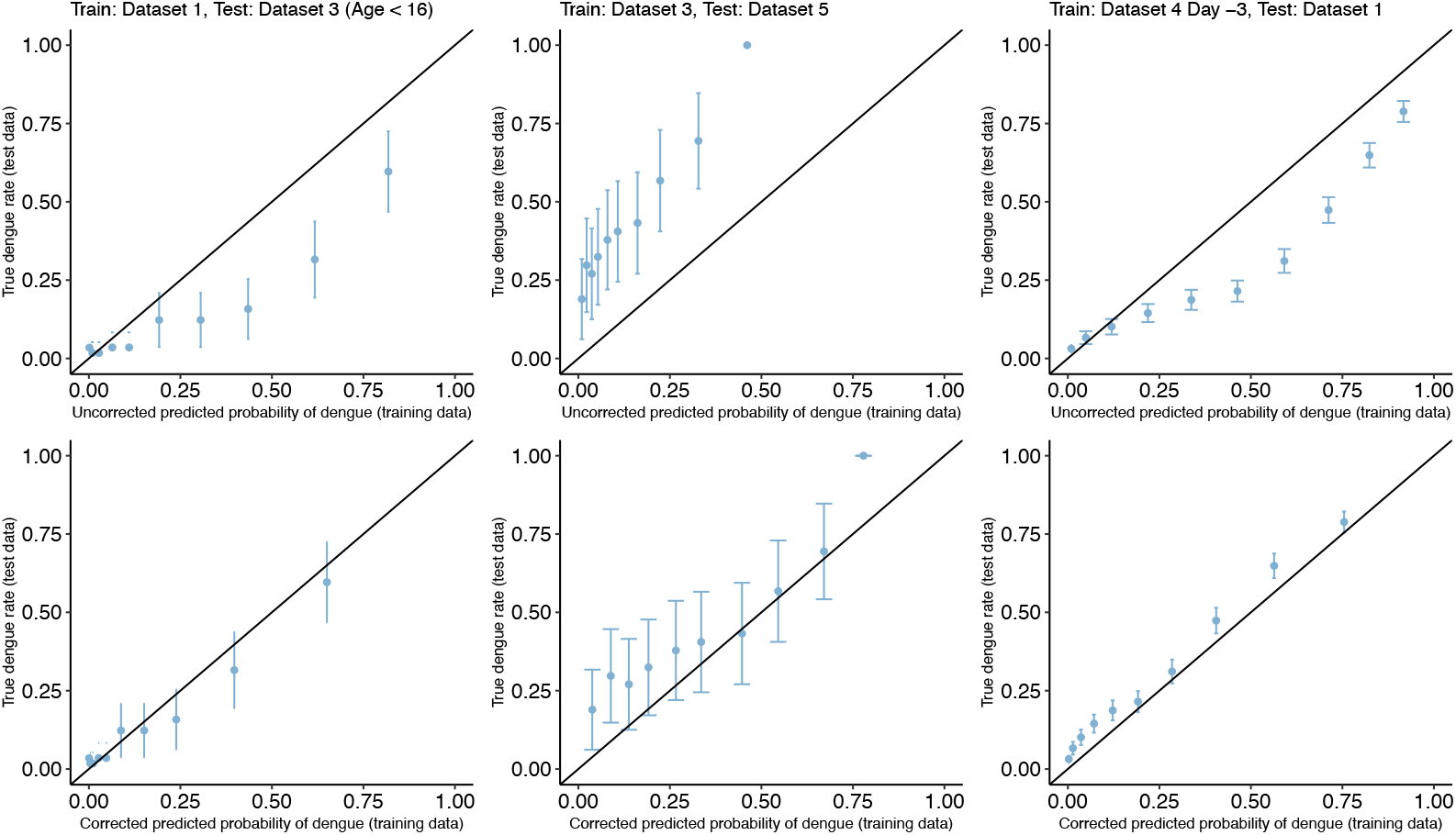
Example calibration plots for several training/test pairs. The logistic regression model is fit on the training set, then applied to the test set to calculate a predicted probability for each test observation. These predicted probabilities are binned into deciles, and the true rate of dengue cases in each bin is plotted against the bin midpoint. Plots are shown for both the raw predictions, and for the label shift-adjusted predictions using Eq. (1).

### Estimating dengue prevalence in test data

The corrected calibration plots in Fig. 4 require the true dengue prevalence in each test set. Fig. 5 shows the results of three different estimates of dengue prevalence in the test data, for each training/test pair. The discretization method (36) performed worst, with several estimates outside the (0, 1) range, and a median log ratio (i.e., log(estimated prevalence*/*true prevalence)) of 0.244 compared to the true dengue prevalence. Performance of the fixed point method (34) was also poor, with a median log ratio of 0.184 compared to the true prevalence. Somewhat surprisingly, simply estimating prevalence with the mean of the uncorrected predicted probabilities performed best, with a median log ratio of 0.016. However, none of the three methods is consistently reliable, with particularly poor performance on test Datasets 4 and 5.

**Fig 5.**
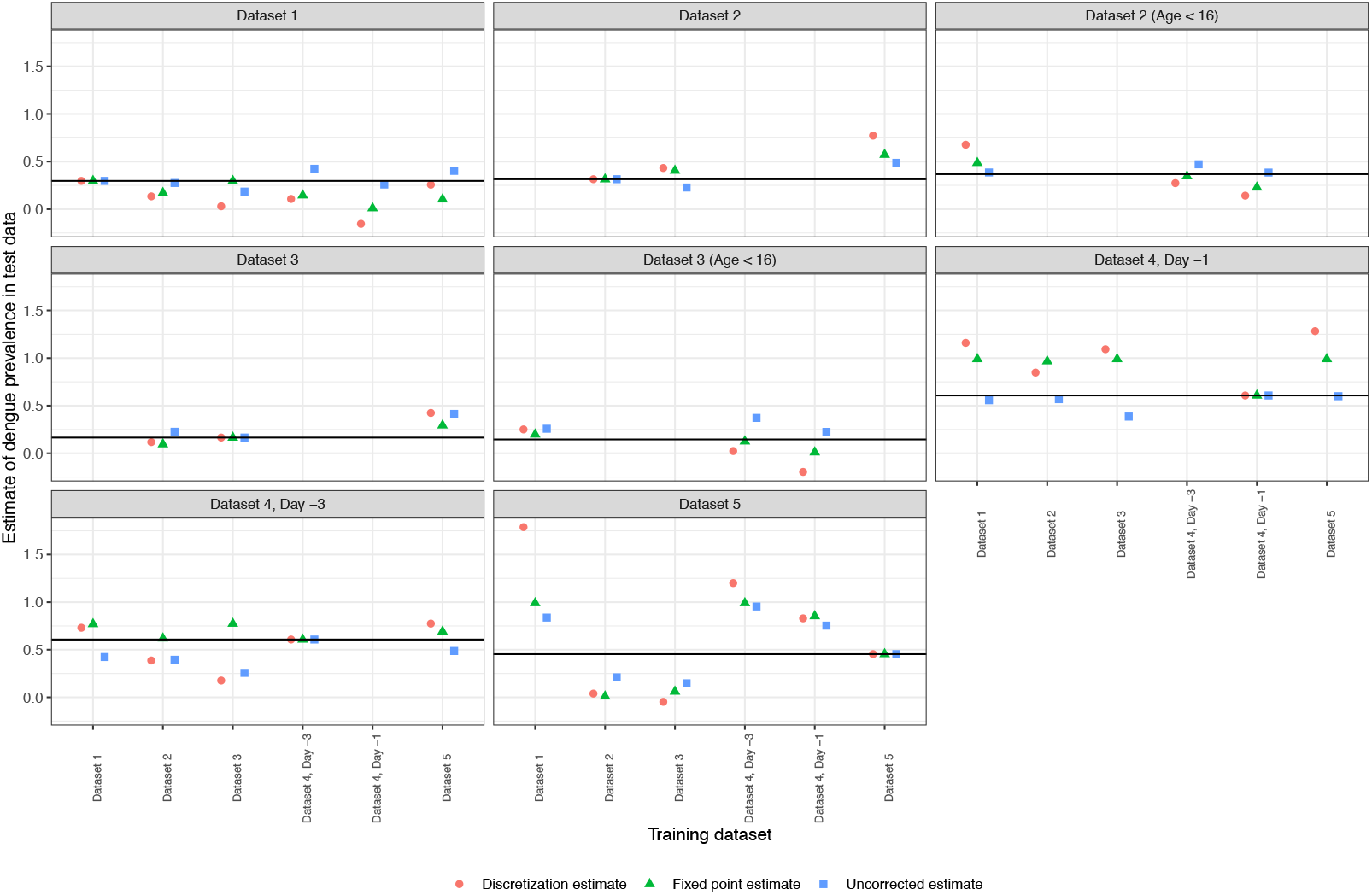
Estimating prevalence of dengue in test data. For each training/test pair, the prevalence of dengue in the test data is estimated by the discretization method, the fixed point method, and by simply averaging the predicted probabilities. The solid horizontal line in each panel represents the true prevalence of dengue in that test set.

### Other explanatory variables

Predictive performance for logistic regression models on each pair of training and test datasets, for the three different alternative sets of explanatory variables (Table 1), are shown in Table S4, Table S5, and Table S6. Overall, performance on the original subset (age, white blood cell count, and platelet count) is comparable or better than performance with other explanatory variables in the model.

For the first alternative subset, AUC is around 0.8 for most logistic regression models considered. An exception occurs when training the full model (all variables in Alternative Subset 1) or the BSS model (the results of best subset selection with AIC) on Dataset 1, and predicting on Dataset 3 or Dataset 5. In these cases, performance on the test data is extremely poor, with an AUC *below* 0.5. This poor performance is not seen with logistic regression on the original explanatory variables.

For the second alternative subset, the difference in predictive performance between the three different logistic regression models (original variables, all variables in Alternative Subset 2, and the BSS model) is very small (generally within 0.01 or 0.02) for all pairs of training and test sets. For the third alternative subset, performance is again similar between all three logistic regression models, particularly when the age range is restricted to be comparable.

These results show that logistic regression with age, white blood cell count, and platelet count achieves similar performance to larger models with additional explanatory variables, which aligns with previous results from Tuan *et al.* (8) and Potts *et al.* (28). Furthermore, in some cases (see Alternative Subset 1 in particular) using fewer variables allowed for improved generalizability to new data.

### Other classification methods

Predictive performance for logistic regression models is compared to decision trees and SVMs, for each set of explanatory variables shown in Table 1, in Fig. S3, Fig. S4, Fig. S5, and Fig. S6.

In general, it appears that the logistic regression model, as a parametric model with a very specific shape assumption, is the most susceptible to extrapolation issues when the datasets have different age ranges. Otherwise, the AUC values for the logistic regression model tend to be higher than for the decision tree, especially when Dataset 1 or Dataset 4 is the test set. The exception is predicting on Dataset 5, for which the decision tree outperforms both the logistic regression and the SVM. The SVM performs similarly to the logistic regression model, with slightly worse performance on Datasets 1 and 2, but less sensitivity to age range differences.

Finally, we note that for all three models, performance on the different sets of explanatory variables (Table 1) is similar, which further supports the choice of age, white blood cell count, and platelet count as our primary explanatory variables of interest.

## Discussion

Many previous studies have investigated the use of statistical models to diagnose dengue in patients. When these models work well, they have the potential to quickly identify dengue fever without requiring potentially expensive or slow laboratory tests. However, each model has been developed for a different population, and collecting training data and fitting an appropriate model can be laborious and time-consuming tasks. If existing models can generalize to new populations, practitioners could make quick diagnoses without needing to create their own new model.

The goal of this paper is to assess the generalizability of a dengue classifier to new populations. We have focused on the Early Dengue Classifier proposed by Tuan *et al.* (8) as a good candidate, because it is easy for practitioners to use, it does not involve too many explanatory variables (and Tuan *et al.* demonstrated similar performance to more complicated models), and the three covariates have been identified as important dengue predictors in multiple other studies. Our analysis explores whether the model performs well when trained on a new dataset (in-sample performance), and whether a pre-trained model can predict well on new test data from a different population (out-of-sample performance). Furthermore, we compare the performance of this model with other logistic regression models incorporating more explanatory variables, and with two other classification methods. We found comparable (or occasionally worse) performance when using larger models, and also when using decision trees and support vector machines. We therefore focus on the simpler logistic regression model proposed by Tuan *et al.* (8) for its ease of use, its interpretability, and its simplicity: requiring few variables makes it easier to implement in new settings.

Our in-sample results in Table 2 show that, when re-trained on data from new settings, this logistic regression model performs well in a variety of settings with different age ranges and dengue prevalences. Indeed, the in-sample AUCs for each of the five datasets are above 0.75, and are between 0.8 and 0.9 for four of the five datasets. The similarities in AUCs across datasets suggest that the chosen logistic regression model is an effective choice for dengue classification in a wide variety of settings, and could be adapted to new environments by collecting new training sets. This performance is also similar to the AUCs reported in many previous studies using a range of classification techniques (8; 23; 26; 27), which supports that the model is competitive with other classifiers.

After confirming that the logistic regression model is a good choice across datasets, we next assessed whether the model could be applied to data from new settings *without* retraining. If the model were generalizable, we would expect its performance on a new dataset to be similar to the in-sample performance we observe. However, there are several potential challenges when applying an existing model to new data: differences in feature distributions (e.g., different age groups), different rates of dengue prevalence, and selecting an appropriate threshold for binary classification.

When we assessed the generalizability of the model with no restriction on the age range of the datasets, the AUC values for the training/test pairs that have a similar or the training dataset has a wider age range were all above 0.75, though most of the AUC values were slightly lower than the in-sample performances, as expected. For training/test pairs with training datasets that have a narrower age range, we observed substantial declines in the AUC values compared with their in-sample performances (Table S2). The disparities in the generalizing performance demonstrate that the model trained on a dataset with a broader age range can extrapolate well when tested on a dataset with a narrower age range but not vice versa.

When we re-assessed the generalizability of the model with restrictions on the age range of the datasets, there were improvements shown in AUC values for all datasets (Fig. 3), which means that having similar age ranges in both training and test datasets can improve the generalizability of the model. Since the model has an AUC value above and is similar to the in-sample performance for all training/testing pairs with similar age ranges, the logistic regression model we have studied in this paper can be applied to a new dataset with minimal reduction in its overall performance (as measured by AUC, which summarizes the overall ability to distinguish dengue and non-dengue cases) if the model is trained on a similar age range with the new dataset.

However, AUC does not capture whether predicted probabilities are calibrated, nor how to choose a threshold when making binary predictions. In the assessment and re-assessment of the generalizability, we observed an imbalance between sensitivity and specificity (Table S2). When we evaluated the model with no restrictions on the age range, all training/test pairs have shown a great imbalance between sensitivity and specificity for both the thresholds from the training and testing datasets. The imbalance is somewhat alleviated when we re-evaluated the model with restricted age ranges using only the threshold from the training datasets. However, the discrepancy between sensitivity and specificity still existed for Dataset 4 Day-1 and Dataset 5 (Fig. 3). Thus, although restricting the age range can mitigate the issue of imbalanced sensitivity and specificity, it is only to an extent. Since having a model with imbalanced sensitivity and specificity may not be helpful for practitioners making diagnoses, one would need to find an alternative way of choosing a threshold for the model to be effective in practice.

Additionally, although the AUC value of the model appears to be relatively good, the predictions were consistently higher or lower than the desired probabilities (Fig. 4) due to the discrepancies in dengue prevalence between the training and testing datasets, which is expected in practice. This suggests that an existing model might not be applicable to new data, without adjusting for differences in dengue prevalence.

The label shift assumption (34; 36) provides a natural method to account for changes in disease prevalence, and the datasets have roughly similar distributions of white blood cell count and platelet count, conditional on dengue status (Fig. 1). Furthermore, in many of the training/test pairs, the predicted probabilities could be calibrated using a label shift assumption (Fig. 4, and Fig. S1 and Fig. S2). However, label shift adjustment requires knowing the dengue prevalence in the test set. If the population prevalence is unknown, or changes (e.g. due to seasonal trends), calibrating the model predictions would be impossible without collecting new data. Attempts to leverage the label shift assumption to estimate test prevalence were unsuccessful (Fig. 5).

In conclusion, we have shown that the logistic regression model from Tuan et al. can be applied to a new dataset without re-training, but its ability to help clinicians make diagnoses is limited. Specifically, for the model to generalize to a new dataset, the age range of the training and test datasets should be roughly the same, or the age range of the training dataset should be wider than the test dataset. Although having the age range restricted can ensure the model’s general predictive ability, it may still not be very suitable to use the model in practice since it is hard to find an appropriate threshold value to generate a relatively balanced sensitivity and specificity. In future studies, to improve the model’s efficiency in clinical settings, one could focus on investigating how to choose a threshold that can result in a balanced sensitivity and specificity. Additionally, since the predicted probabilities made by the model tend to be biased, and the technique required to calibrate it might be too complicated to exert in practice, one could also focus on improving the model’s performance by adjusting the distribution of the explanatory variables before training or the model so that its predictive probabilities match with the desired probabilities without calibration.

## Supporting information

Table S1. Summary statistics for each dataset.

Table S2. Full performance metrics for each combination of training/test datasets.

Table S3. Label shift adjustment and estimation for each training/test pair.

Table S4. In-sample and generalizability performance metrics for logistic regression with each training/test pair, Alternative Subset 1.

Table S5. In-sample and generalizability performance metrics for logistic regression with each training/test pair, Alternative Subset 2.

Table S6. In-sample and generalizability performance metrics for logistic regression with each training/test pair, Alternative Subset 3.

Fig. S1 Calibration plots for all training/test pairs (uncorrected predictions). The logistic regression model is fit on the training set, and applied to the test set to calculate predicted probabilities. These raw predicted probabilities are compared against true instances of dengue.

Fig. S2 Calibration plots for all training/test pairs (label shift-adjusted predictions). The logistic regression model is fit on the training set, and applied to the test set to calculate predicted probabilities. The predicted probabilities are adjusted with a label shift correction to account for a different rate of dengue fever in the test dataset (see Eq. 1 in the manuscript).

Fig. S3 Comparison of predictive performance with the original explanatory variables for three different classification methods, for each pair of training/test datasets.

Fig. S4 Comparison of predictive performance with the alternative subset 1 explanatory variables for three different classification methods, for each pair of training/test datasets.

Fig. S5 Comparison of predictive performance with the alternative subset 2 explanatory variables for three different classification methods, for each pair of training/test datasets.

Fig. S6 Comparison of predictive performance with the alternative subset 3 explanatory variables for three different classification methods, for each pair of training/test datasets.

## Supplementary Material

June 17, 2024

**Supplementary Fig 1.**
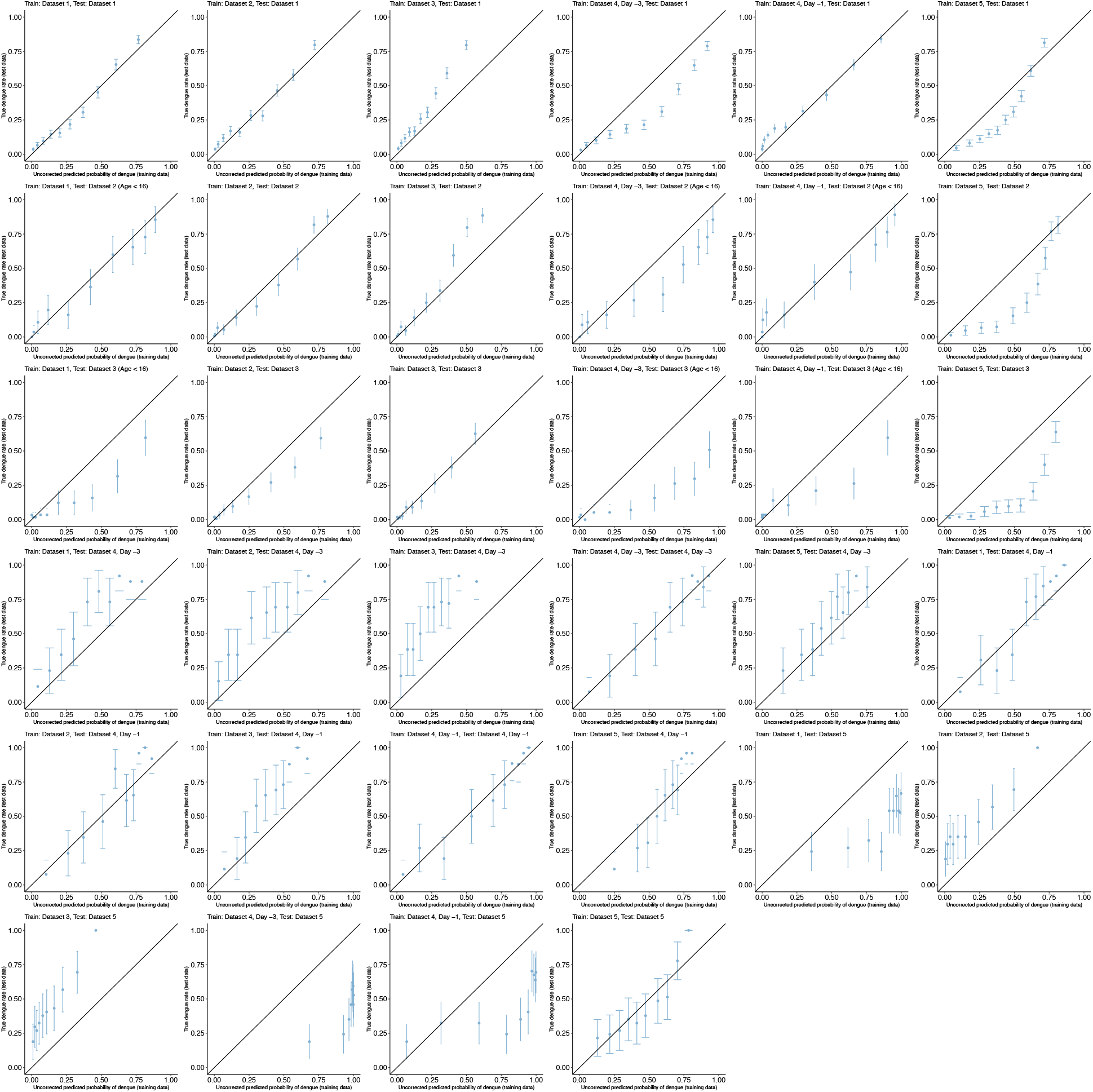
Calibration plots for all training/test pairs (uncorrected predictions). The logistic regression model is fit on the training set, and applied to the test set to calculate predicted probabilities. These raw predicted probabilities are compared against true instances of dengue.

**Supplementary Fig 2.**
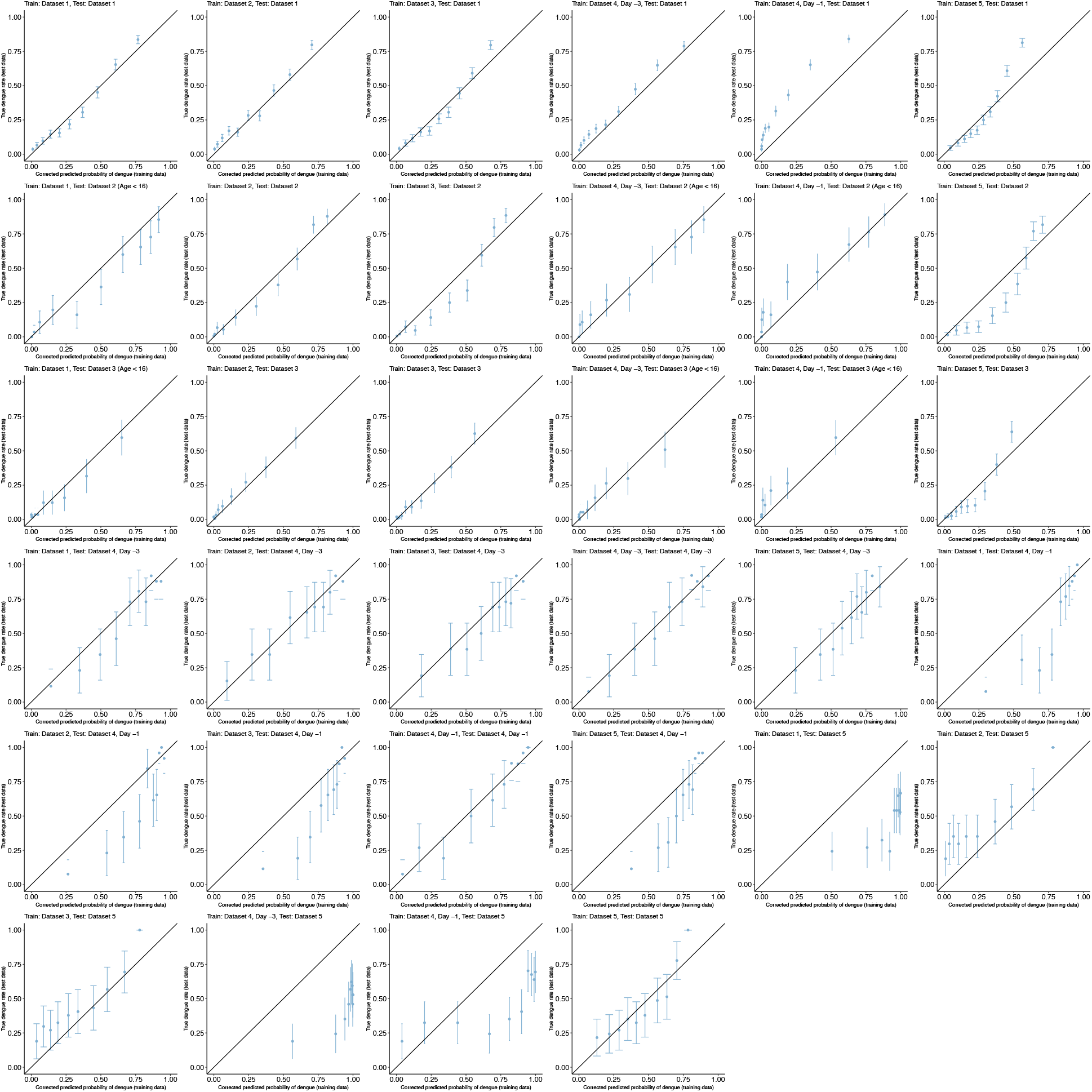
Calibration plots for all training/test pairs (label shift-adjusted predictions). The logistic regression model is fit on the training set, and applied to the test set to calculate predicted probabilities. The predicted probabilities are adjusted with a label shift correction to account for a different rate of dengue fever in the test dataset (see Eq. 1 in the manuscript).

**Supplementary Fig 3.**
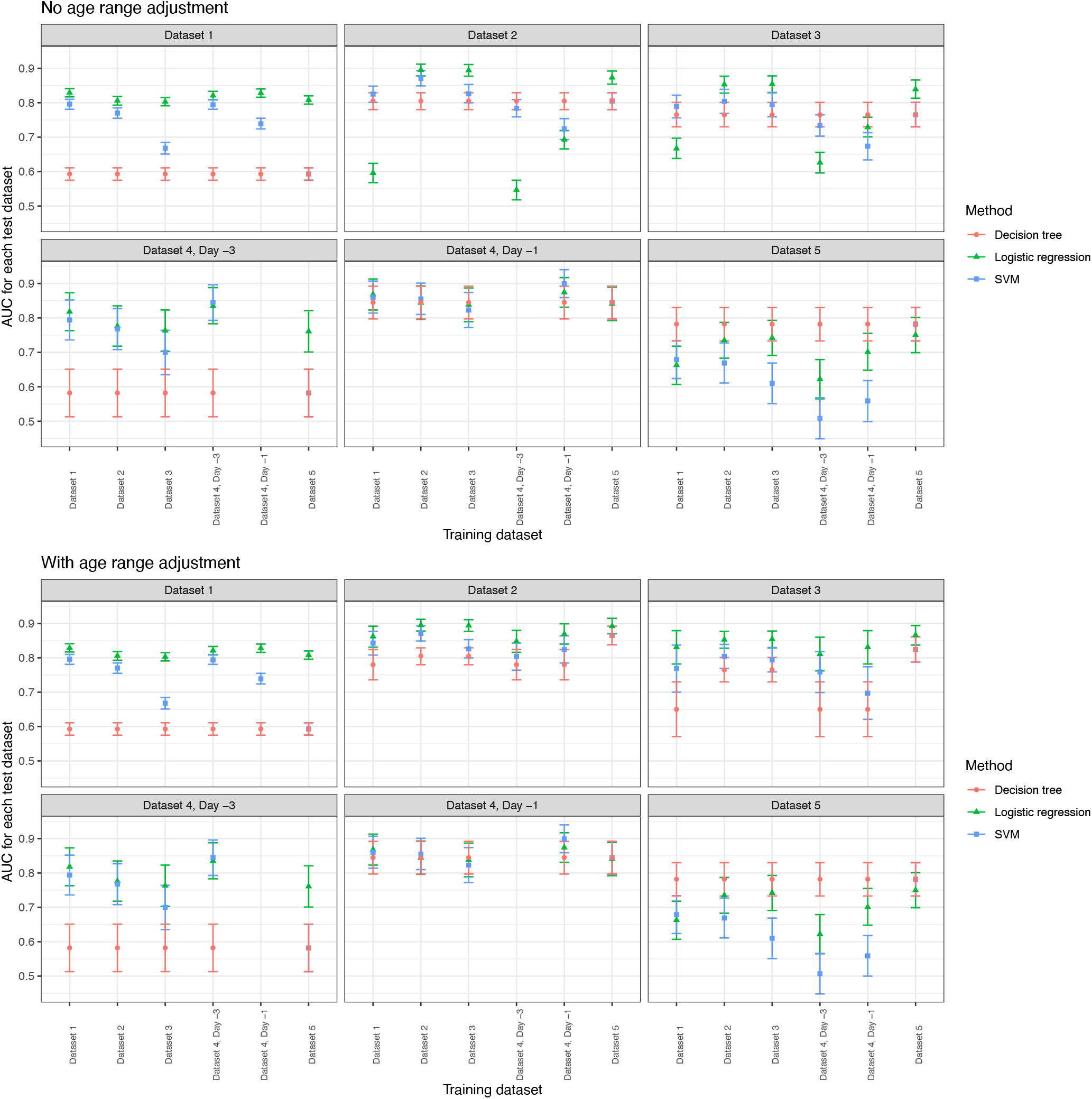
Comparison of predictive performance with the original explanatory variables for three different classification methods, for each pair of training/test datasets. Recall that the original explanatory variables are Age, WBC, and PLT (see Table 1 in the main manuscript). Each panel represents one test set, with the predictive performance on that test set (the AUC) displayed for each training set. Performance is shown with and without age range restrictions. Without age range restrictions, all complete cases in the training and test datasets are used. With age range restrictions, the test dataset is restricted (if possible) to match the age range of the training data (< 16 if training data is Dataset 1 or Dataset 4 and test data is Dataset 2 or Dataset 3; > 16 if training data is Dataset 5 and test data is Dataset 2 or Dataset 3).

**Supplementary Fig 4.**
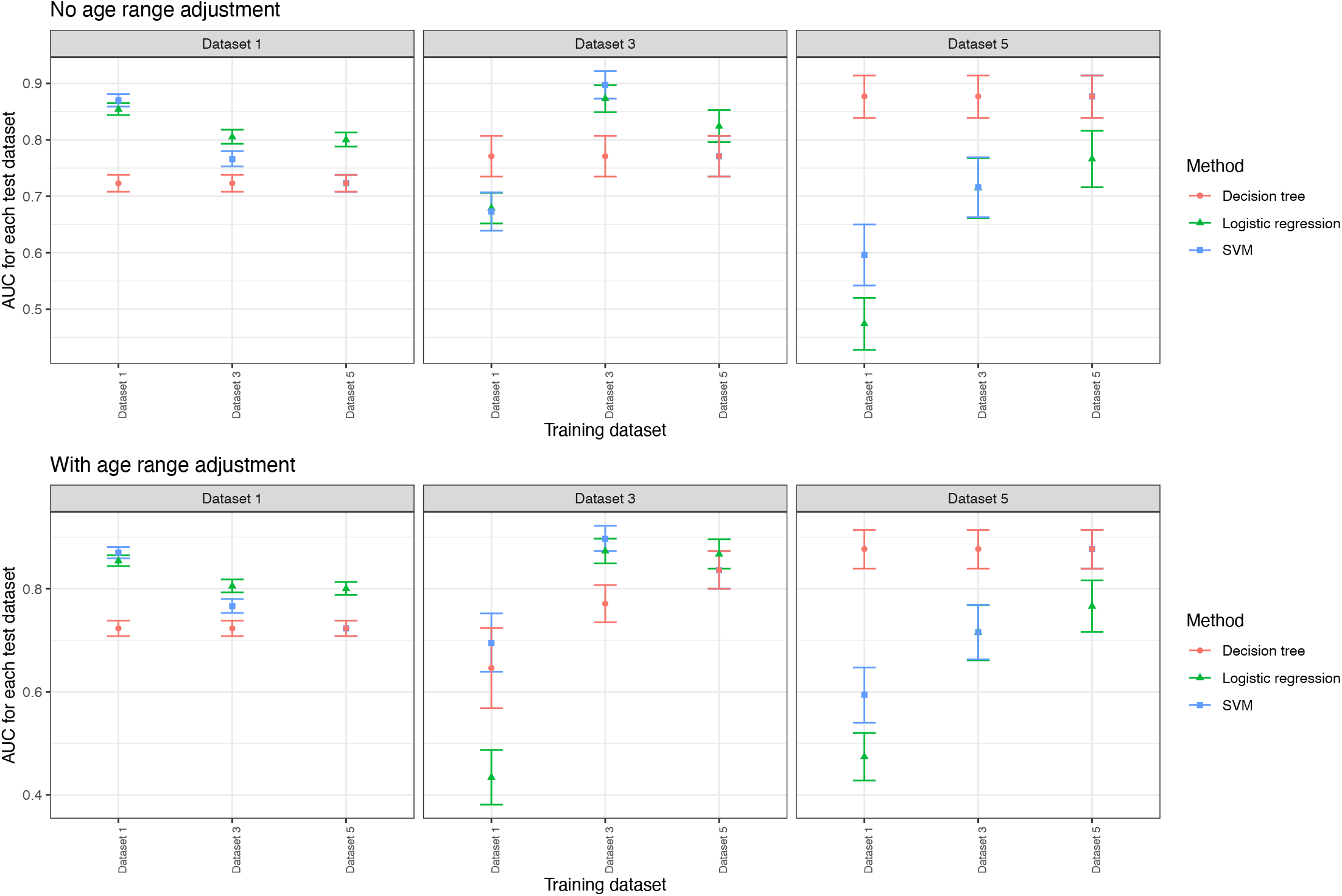
Comparison of predictive performance with the alternative subset 1 explanatory variables for three different classification methods, for each pair of training/test datasets. The variables in Alternative Subset 1 can be found in Table 1. Each panel represents one test set, with the predictive performance on that test set (the AUC) displayed for each training set. Performance is shown with and without age range restrictions. Without age range restrictions, all complete cases in the training and test datasets are used. With age range restrictions, the test dataset is restricted (if possible) to match the age range of the training data (< 16 if training data is Dataset 1 or Dataset 4 and test data is Dataset 2 or Dataset 3; > 16 if training data is Dataset 5 and test data is Dataset 2 or Dataset 3).

**Supplementary Fig 5.**
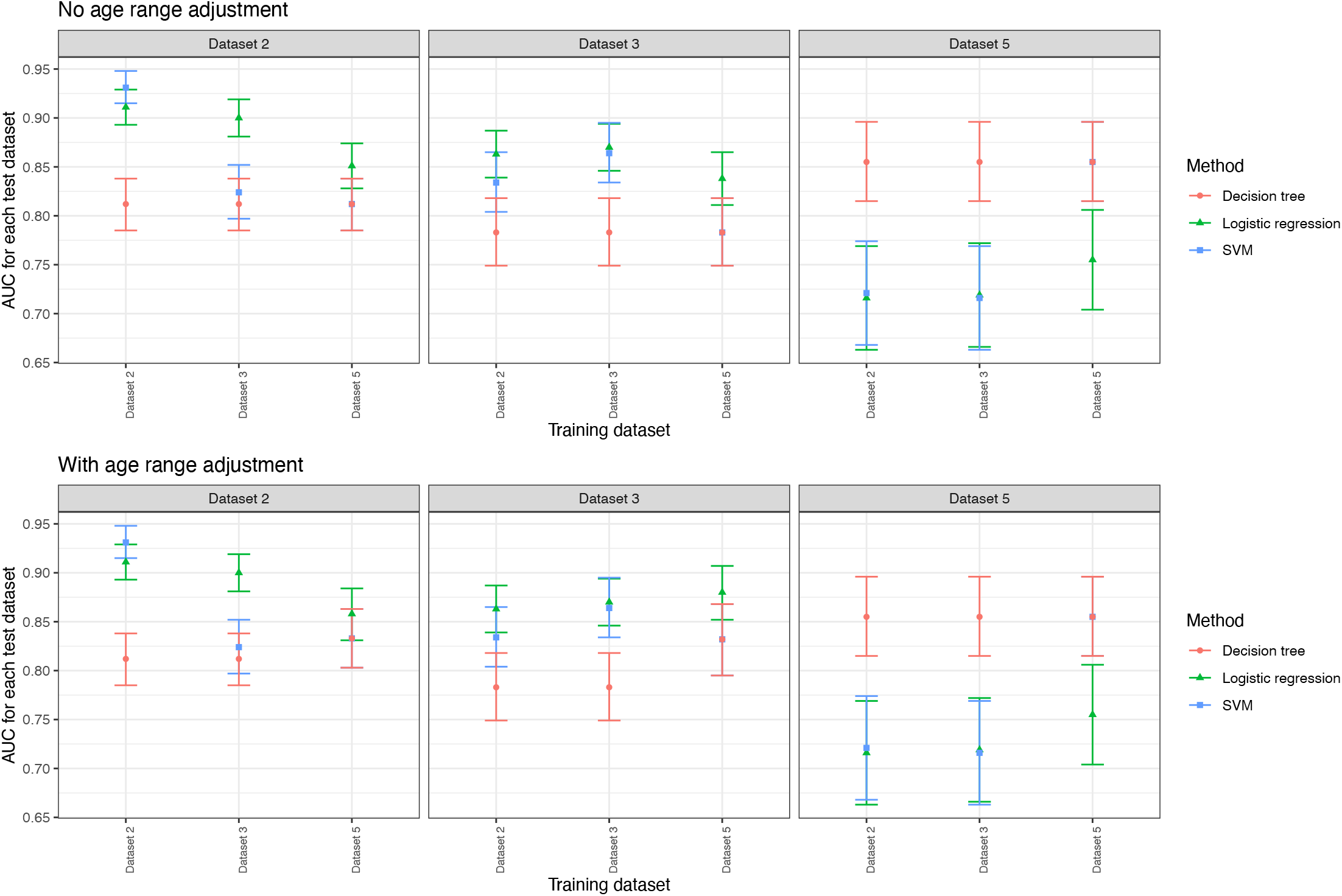
Comparison of predictive performance with the alternative subset 2 explanatory variables for three different classification methods, for each pair of training/test datasets. The variables in Alternative Subset 2 can be found in Table 1. Each panel represents one test set, with the predictive performance on that test set (the AUC) displayed for each training set. Performance is shown with and without age range restrictions. Without age range restrictions, all complete cases in the training and test datasets are used. With age range restrictions, the test dataset is restricted (if possible) to match the age range of the training data (< 16 if training data is Dataset 1 or Dataset 4 and test data is Dataset 2 or Dataset 3; > 16 if training data is Dataset 5 and test data is Dataset 2 or Dataset 3).

**Supplementary Fig 6.**
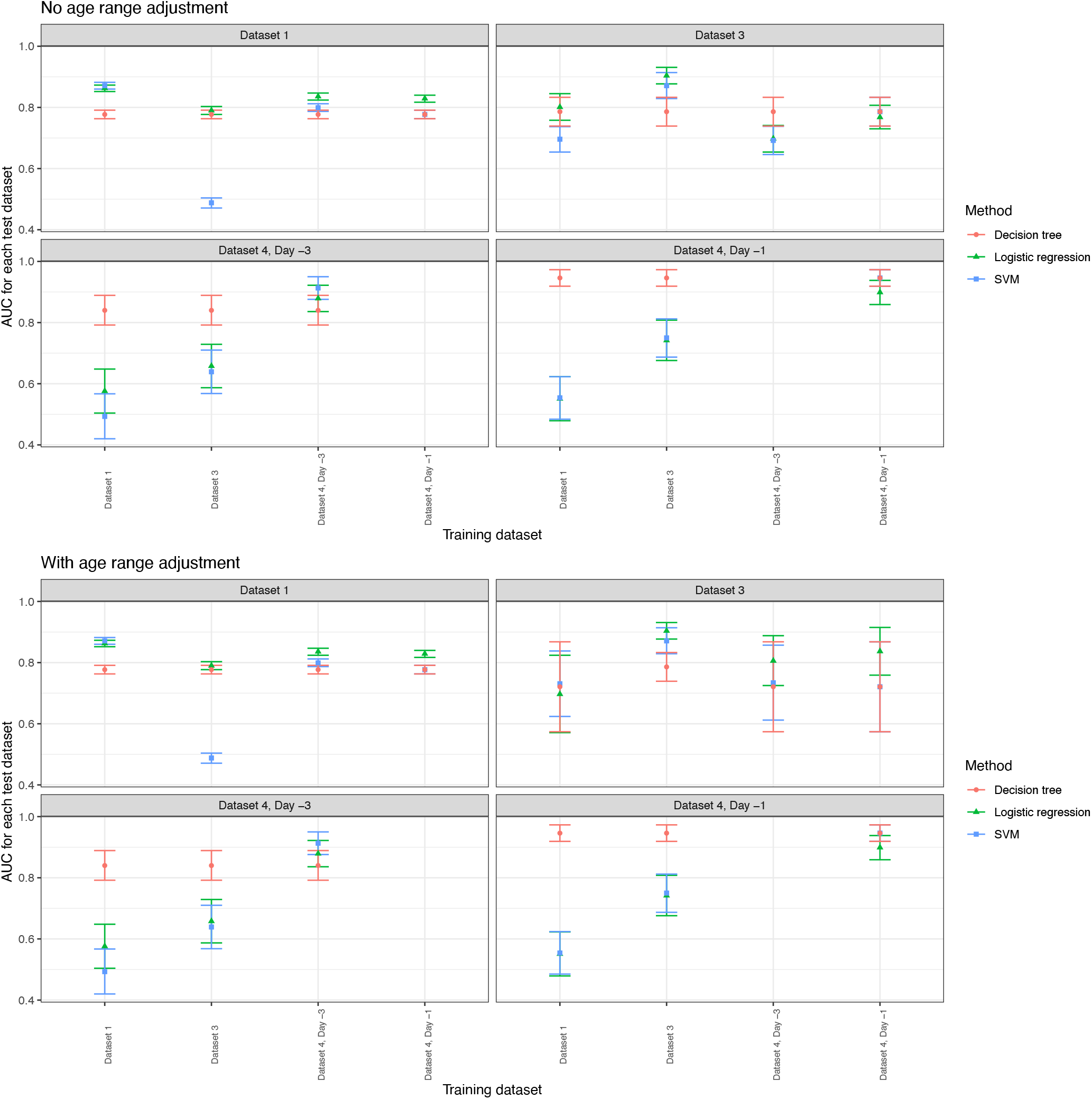
Comparison of predictive performance with the alternative subset 3 explanatory variables for three different classification methods, for each pair of training/test datasets. The variables in Alternative Subset 3 can be found in Table 1. Each panel represents one test set, with the predictive performance on that test set (the AUC) displayed for each training set. Performance is shown with and without age range restrictions. Without age range restrictions, all complete cases in the training and test datasets are used. With age range restrictions, the test dataset is restricted (if possible) to match the age range of the training data (< 16 if training data is Dataset 1 or Dataset 4 and test data is Dataset 2 or Dataset 3; > 16 if training data is Dataset 5 and test data is Dataset 2 or Dataset 3).

**Supplementary Table 1.** Summary statistics for each dataset.

|  | Dataset 1 (N=5724) | Dataset 2 (N=1485) | Dataset 3 (N=1552) | Dataset 4, Day -1 (N=257) | Dataset 4, Day -3 (N=257) | Dataset 5 (N=368) |
| --- | --- | --- | --- | --- | --- | --- |
| **Dengue** |  |  |  |  |  |  |
| Negative | 4026 (70.3%) | 1018 (68.6%) | 1296 (83.5%) | 101 (39.3%) | 101 (39.3%) | 201 (54.6%) |
| Positive | 1698 (29.7%) | 467 (31.4%) | 256 (16.5%) | 156 (60.7%) | 156 (60.7%) | 167 (45.4%) |
| **Age** |  |  |  |  |  |  |
| Mean (SD) | 6.8 (3.6) | 25.6 (19.7) | 23.6 (17.2) | 7.7 (3.1) | 7.7 (3.1) | 33.0 (11.6) |
| Median | 6.0 (4.0, 9.0) | 20.8 (9.1, 37.9) | 20.0 (10.0, 33.0) | 8.5 (4.5, 10.5) | 8.5 (4.5, 10.5) | 30.2 (23.8, 39.9) |
| Range | 1.0 - 15.0 | 1.0 - 98.0 | 1.0 - 82.0 | 0.5 - 14.5 | 0.5 - 14.5 | 17.0 - 78.4 |
| **White Blood Cell Count (WBC)** |  |  |  |  |  |  |
| Mean (SD) | 8.6 (4.8) | 9.0 (7.0) | 10.3 (6.9) | 4.2 (2.6) | 6.1 (3.9) | 7.3 (3.7) |
| Median | 7.5 (5.0, 11.0) | 6.9 (4.0, 12.2) | 8.7 (5.3, 13.2) | 3.4 (2.4, 5.1) | 5.1 (3.5, 7.8) | 6.7 (4.6, 9.3) |
| Range | 0.9 - 49.6 | 0.5 - 70.5 | 0.2 - 47.3 | 0.6 - 17.9 | 1.4 - 35.8 | 1.4 - 21.1 |
| **Platelet Count (PLT)** |  |  |  |  |  |  |
| Mean (SD) | 231.5 (80.0) | 187.0 (123.4) | 215.4 (138.8) | 173.5 (81.2) | 215.6 (80.8) | 222.5 (98.6) |
| Median | 224.0 (179.0, 277.0) | 160.0 (92.0, 253.0) | 194.0 (108.8, 294.0) | 166.0 (116.0, 222.0) | 205.0 (156.0, 262.0) | 222.5 (153.8, 282.8) |
| Range | 18.0 - 829.0 | 7.0 - 855.0 | 0.0 - 862.0 | 18.0 - 427.0 | 11.0 - 464.0 | 0.0 - 740.0 |

**Supplementary Table 2.** Full performance metrics for each combination of training/test datasets.

| Training data | Test data | Threshold | Sensitivity | Specificity | PPV | NPV | AUC |
| --- | --- | --- | --- | --- | --- | --- | --- |
| Dataset 1 | Dataset 1 | Training threshold | 0.747 (0.725, 0.767) | 0.763 (0.75, 0.776) | 0.571 (0.55, 0.591) | 0.877 (0.866, 0.888) | 0.829 (0.817, 0.841) |
| Dataset 1 | Dataset 2 | Training threshold | 0.938 (0.912, 0.958) | 0.296 (0.268, 0.325) | 0.379 (0.351, 0.408) | 0.912 (0.876, 0.94) | 0.596 (0.568, 0.624) |
| Dataset 1 | Dataset 2 | Test threshold | 0.899 (0.868, 0.925) | 0.344 (0.315, 0.374) | 0.386 (0.357, 0.416) | 0.882 (0.846, 0.912) | 0.596 (0.568, 0.624) |
| Dataset 1 | Dataset 3 | Training threshold | 0.914 (0.873, 0.945) | 0.366 (0.339, 0.393) | 0.222 (0.197, 0.248) | 0.956 (0.934, 0.972) | 0.667 (0.638, 0.697) |
| Dataset 1 | Dataset 3 | Test threshold | 0.965 (0.934, 0.984) | 0.297 (0.272, 0.323) | 0.213 (0.19, 0.238) | 0.977 (0.957, 0.99) | 0.667 (0.638, 0.697) |
| Dataset 1 | Dataset 4, Day -3 | Training threshold | 0.827 (0.758, 0.883) | 0.693 (0.593, 0.781) | 0.806 (0.736, 0.864) | 0.722 (0.621, 0.808) | 0.818 (0.763, 0.873) |
| Dataset 1 | Dataset 4, Day -3 | Test threshold | 0.449 (0.369, 0.53) | 0.921 (0.85, 0.965) | 0.897 (0.808, 0.955) | 0.52 (0.444, 0.595) | 0.818 (0.763, 0.873) |
| Dataset 1 | Dataset 4, Day -1 | Training threshold | 0.936 (0.885, 0.969) | 0.446 (0.347, 0.548) | 0.723 (0.656, 0.783) | 0.818 (0.691, 0.909) | 0.868 (0.823, 0.913) |
| Dataset 1 | Dataset 4, Day -1 | Test threshold | 0.705 (0.627, 0.775) | 0.881 (0.802, 0.937) | 0.902 (0.834, 0.948) | 0.659 (0.573, 0.739) | 0.868 (0.823, 0.913) |
| Dataset 1 | Dataset 5 | Training threshold | 0.994 (0.967, 1) | 0.06 (0.031, 0.102) | 0.468 (0.415, 0.521) | 0.923 (0.64, 0.998) | 0.663 (0.607, 0.718) |
| Dataset 1 | Dataset 5 | Test threshold | 0.97 (0.932, 0.99) | 0.095 (0.058, 0.144) | 0.471 (0.417, 0.525) | 0.792 (0.578, 0.929) | 0.663 (0.607, 0.718) |
| Dataset 2 | Dataset 1 | Training threshold | 0.554 (0.53, 0.578) | 0.879 (0.868, 0.889) | 0.659 (0.633, 0.683) | 0.824 (0.812, 0.835) | 0.806 (0.793, 0.818) |
| Dataset 2 | Dataset 1 | Test threshold | 0.689 (0.666, 0.711) | 0.765 (0.752, 0.778) | 0.553 (0.531, 0.574) | 0.854 (0.842, 0.865) | 0.806 (0.793, 0.818) |
| Dataset 2 | Dataset 2 | Training threshold | 0.805 (0.766, 0.84) | 0.847 (0.823, 0.868) | 0.707 (0.666, 0.745) | 0.905 (0.884, 0.922) | 0.895 (0.878, 0.912) |
| Dataset 2 | Dataset 3 | Training threshold | 0.625 (0.563, 0.685) | 0.856 (0.835, 0.874) | 0.461 (0.408, 0.515) | 0.92 (0.904, 0.935) | 0.853 (0.828, 0.877) |
| Dataset 2 | Dataset 3 | Test threshold | 0.84 (0.789, 0.883) | 0.715 (0.69, 0.74) | 0.368 (0.329, 0.409) | 0.958 (0.943, 0.969) | 0.853 (0.828, 0.877) |
| Dataset 2 | Dataset 4, Day -3 | Training threshold | 0.571 (0.489, 0.649) | 0.802 (0.711, 0.875) | 0.817 (0.731, 0.884) | 0.547 (0.463, 0.629) | 0.776 (0.718, 0.835) |
| Dataset 2 | Dataset 4, Day -3 | Test threshold | 0.378 (0.302, 0.459) | 0.911 (0.838, 0.958) | 0.868 (0.764, 0.938) | 0.487 (0.414, 0.56) | 0.776 (0.718, 0.835) |
| Dataset 2 | Dataset 4, Day -1 | Training threshold | 0.891 (0.831, 0.935) | 0.624 (0.522, 0.718) | 0.785 (0.717, 0.843) | 0.787 (0.682, 0.871) | 0.844 (0.796, 0.893) |
| Dataset 2 | Dataset 4, Day -1 | Test threshold | 0.667 (0.587, 0.74) | 0.782 (0.689, 0.858) | 0.825 (0.748, 0.887) | 0.603 (0.514, 0.687) | 0.844 (0.796, 0.893) |
| Dataset 2 | Dataset 5 | Training threshold | 0.335 (0.264, 0.412) | 0.96 (0.923, 0.983) | 0.875 (0.768, 0.944) | 0.635 (0.578, 0.689) | 0.735 (0.683, 0.787) |
| Dataset 2 | Dataset 5 | Test threshold | 0.347 (0.275, 0.425) | 0.945 (0.904, 0.972) | 0.841 (0.733, 0.918) | 0.635 (0.578, 0.69) | 0.735 (0.683, 0.787) |
| Dataset 3 | Dataset 1 | Training threshold | 0.689 (0.666, 0.711) | 0.767 (0.753, 0.78) | 0.555 (0.533, 0.576) | 0.854 (0.842, 0.865) | 0.803 (0.791, 0.815) |
| Dataset 3 | Dataset 1 | Test threshold | 0.438 (0.414, 0.462) | 0.923 (0.914, 0.931) | 0.706 (0.677, 0.733) | 0.796 (0.784, 0.807) | 0.803 (0.791, 0.815) |
| Dataset 3 | Dataset 2 | Training threshold | 0.876 (0.842, 0.904) | 0.744 (0.716, 0.77) | 0.61 (0.572, 0.648) | 0.929 (0.909, 0.946) | 0.894 (0.877, 0.911) |
| Dataset 3 | Dataset 2 | Test threshold | 0.548 (0.502, 0.594) | 0.951 (0.936, 0.963) | 0.837 (0.79, 0.876) | 0.821 (0.798, 0.843) | 0.894 (0.877, 0.911) |
| Dataset 3 | Dataset 3 | Training threshold | 0.781 (0.726, 0.83) | 0.78 (0.757, 0.802) | 0.412 (0.368, 0.458) | 0.948 (0.932, 0.96) | 0.854 (0.829, 0.878) |
| Dataset 3 | Dataset 4, Day -3 | Training threshold | 0.731 (0.654, 0.799) | 0.663 (0.562, 0.754) | 0.77 (0.694, 0.835) | 0.615 (0.517, 0.706) | 0.763 (0.703, 0.823) |
| Dataset 3 | Dataset 4, Day -3 | Test threshold | 0.038 (0.014, 0.082) | 0.98 (0.93, 0.998) | 0.75 (0.349, 0.968) | 0.398 (0.336, 0.461) | 0.763 (0.703, 0.823) |
| Dataset 3 | Dataset 4, Day -1 | Training threshold | 0.923 (0.869, 0.96) | 0.485 (0.384, 0.587) | 0.735 (0.667, 0.795) | 0.803 (0.682, 0.894) | 0.838 (0.789, 0.887) |
| Dataset 3 | Dataset 4, Day -1 | Test threshold | 0.128 (0.08, 0.191) | 0.99 (0.946, 1) | 0.952 (0.762, 0.999) | 0.424 (0.36, 0.49) | 0.838 (0.789, 0.887) |
| Dataset 3 | Dataset 5 | Training threshold | 0.425 (0.349, 0.504) | 0.886 (0.833, 0.926) | 0.755 (0.656, 0.838) | 0.65 (0.59, 0.706) | 0.742 (0.691, 0.793) |
| Dataset 3 | Dataset 5 | Test threshold | 0.114 (0.07, 0.172) | 1 (0.982, 1) | 1 (0.824, 1) | 0.576 (0.522, 0.628) | 0.742 (0.691, 0.793) |
| Dataset 4, Day -3 | Dataset 1 | Training threshold | 0.697 (0.675, 0.719) | 0.793 (0.781, 0.806) | 0.587 (0.565, 0.609) | 0.861 (0.85, 0.872) | 0.821 (0.809, 0.833) |
| Dataset 4, Day -3 | Dataset 1 | Test threshold | 0.856 (0.838, 0.872) | 0.574 (0.558, 0.589) | 0.458 (0.441, 0.476) | 0.904 (0.892, 0.915) | 0.821 (0.809, 0.833) |
| Dataset 4, Day -3 | Dataset 2 | Training threshold | 0.901 (0.871, 0.927) | 0.303 (0.274, 0.332) | 0.372 (0.344, 0.401) | 0.87 (0.831, 0.903) | 0.547 (0.518, 0.575) |
| Dataset 4, Day -3 | Dataset 2 | Test threshold | 0.936 (0.91, 0.956) | 0.266 (0.239, 0.295) | 0.369 (0.342, 0.397) | 0.9 (0.861, 0.932) | 0.547 (0.518, 0.575) |
| Dataset 4, Day -3 | Dataset 3 | Training threshold | 0.906 (0.864, 0.939) | 0.363 (0.337, 0.39) | 0.219 (0.195, 0.246) | 0.952 (0.929, 0.969) | 0.626 (0.596, 0.656) |
| Dataset 4, Day -3 | Dataset 3 | Test threshold | 0.969 (0.939, 0.986) | 0.234 (0.211, 0.258) | 0.2 (0.178, 0.223) | 0.974 (0.95, 0.989) | 0.626 (0.596, 0.656) |
| Dataset 4, Day -3 | Dataset 4, Day -3 | Training threshold | 0.833 (0.765, 0.888) | 0.723 (0.625, 0.807) | 0.823 (0.754, 0.879) | 0.737 (0.639, 0.821) | 0.835 (0.783, 0.888) |
| Dataset 4, Day -3 | Dataset 5 | Training threshold | 0.994 (0.967, 1) | 0.045 (0.021, 0.083) | 0.464 (0.411, 0.517) | 0.9 (0.555, 0.997) | 0.622 (0.565, 0.679) |
| Dataset 4, Day -3 | Dataset 5 | Test threshold | 1 (0.978, 1) | 0.03 (0.011, 0.064) | 0.461 (0.409, 0.514) | 1 (0.541, 1) | 0.622 (0.565, 0.679) |
| Dataset 4, Day -1 | Dataset 1 | Training threshold | 0.423 (0.4, 0.447) | 0.951 (0.944, 0.958) | 0.785 (0.757, 0.811) | 0.796 (0.785, 0.808) | 0.828 (0.816, 0.84) |
| Dataset 4, Day -1 | Dataset 1 | Test threshold | 0.676 (0.653, 0.698) | 0.824 (0.812, 0.835) | 0.618 (0.595, 0.64) | 0.858 (0.846, 0.868) | 0.828 (0.816, 0.84) |
| Dataset 4, Day -1 | Dataset 2 | Training threshold | 0.861 (0.826, 0.891) | 0.509 (0.478, 0.54) | 0.446 (0.413, 0.479) | 0.889 (0.86, 0.913) | 0.693 (0.666, 0.719) |
| Dataset 4, Day -1 | Dataset 2 | Test threshold | 0.895 (0.864, 0.921) | 0.453 (0.422, 0.484) | 0.429 (0.397, 0.46) | 0.904 (0.875, 0.928) | 0.693 (0.666, 0.719) |
| Dataset 4, Day -1 | Dataset 3 | Training threshold | 0.781 (0.726, 0.83) | 0.607 (0.58, 0.634) | 0.282 (0.249, 0.317) | 0.934 (0.915, 0.949) | 0.729 (0.701, 0.758) |
| Dataset 4, Day -1 | Dataset 3 | Test threshold | 0.906 (0.864, 0.939) | 0.448 (0.421, 0.476) | 0.245 (0.218, 0.274) | 0.96 (0.942, 0.974) | 0.729 (0.701, 0.758) |
| Dataset 4, Day -1 | Dataset 4, Day -1 | Training threshold | 0.821 (0.751, 0.877) | 0.762 (0.667, 0.841) | 0.842 (0.774, 0.896) | 0.733 (0.638, 0.815) | 0.874 (0.831, 0.917) |
| Dataset 4, Day -1 | Dataset 5 | Training threshold | 0.838 (0.774, 0.891) | 0.368 (0.301, 0.439) | 0.524 (0.463, 0.586) | 0.733 (0.635, 0.816) | 0.701 (0.648, 0.755) |
| Dataset 4, Day -1 | Dataset 5 | Test threshold | 0.892 (0.835, 0.935) | 0.269 (0.209, 0.336) | 0.503 (0.445, 0.562) | 0.75 (0.634, 0.845) | 0.701 (0.648, 0.755) |
| Dataset 5 | Dataset 1 | Training threshold | 0.783 (0.762, 0.802) | 0.67 (0.656, 0.685) | 0.5 (0.481, 0.52) | 0.88 (0.868, 0.891) | 0.808 (0.796, 0.82) |
| Dataset 5 | Dataset 1 | Test threshold | 0.889 (0.873, 0.904) | 0.479 (0.464, 0.495) | 0.419 (0.402, 0.435) | 0.911 (0.898, 0.923) | 0.808 (0.796, 0.82) |
| Dataset 5 | Dataset 2 | Training threshold | 0.936 (0.91, 0.956) | 0.547 (0.516, 0.578) | 0.487 (0.453, 0.52) | 0.949 (0.928, 0.965) | 0.873 (0.854, 0.892) |
| Dataset 5 | Dataset 2 | Test threshold | 0.936 (0.91, 0.956) | 0.578 (0.547, 0.608) | 0.504 (0.47, 0.538) | 0.951 (0.931, 0.967) | 0.873 (0.854, 0.892) |
| Dataset 5 | Dataset 3 | Training threshold | 0.871 (0.824, 0.91) | 0.593 (0.566, 0.62) | 0.297 (0.265, 0.331) | 0.959 (0.943, 0.972) | 0.839 (0.813, 0.866) |
| Dataset 5 | Dataset 3 | Test threshold | 0.969 (0.939, 0.986) | 0.325 (0.299, 0.351) | 0.221 (0.197, 0.246) | 0.981 (0.964, 0.992) | 0.839 (0.813, 0.866) |
| Dataset 5 | Dataset 4, Day -3 | Training threshold | 0.788 (0.716, 0.85) | 0.594 (0.492, 0.691) | 0.75 (0.677, 0.814) | 0.645 (0.539, 0.742) | 0.761 (0.701, 0.821) |
| Dataset 5 | Dataset 4, Day -3 | Test threshold | 0.442 (0.363, 0.524) | 0.881 (0.802, 0.937) | 0.852 (0.756, 0.921) | 0.506 (0.429, 0.582) | 0.761 (0.701, 0.821) |
| Dataset 5 | Dataset 4, Day -1 | Training threshold | 0.949 (0.901, 0.978) | 0.347 (0.255, 0.448) | 0.692 (0.625, 0.753) | 0.814 (0.666, 0.916) | 0.841 (0.792, 0.889) |
| Dataset 5 | Dataset 4, Day -1 | Test threshold | 0.692 (0.614, 0.764) | 0.802 (0.711, 0.875) | 0.844 (0.769, 0.902) | 0.628 (0.538, 0.711) | 0.841 (0.792, 0.889) |
| Dataset 5 | Dataset 5 | Training threshold | 0.707 (0.631, 0.774) | 0.632 (0.561, 0.699) | 0.615 (0.542, 0.684) | 0.722 (0.649, 0.786) | 0.75 (0.699, 0.801) |

**Supplementary Table 3.** Label shift adjustment and estimation for each training/test pair.

| Training data | Test data | Deviance<br>(uncorrected<br>predictions) | Deviance<br>(corrected<br>predictions) | Training<br>prevalence | Test<br>prevalence | Est. test<br>prevalence,<br>average | Est. test<br>prevalence,<br>discretization | Est. test<br>prevalence,<br>fixed point |
| --- | --- | --- | --- | --- | --- | --- | --- | --- |
| Dataset 1 | Dataset 2 (Age < 16) | 0.90 | 0.93 | 0.30 | 0.37 | 0.39 | 0.49 | 0.68 |
| Dataset 1 | Dataset 3 (Age < 16) | 0.78 | 0.67 | 0.30 | 0.14 | 0.26 | 0.20 | 0.25 |
| Dataset 1 | Dataset 4, Day -1 | 0.95 | 1.12 | 0.30 | 0.61 | 0.56 | 0.99 | 1.16 |
| Dataset 1 | Dataset 4, Day -3 | 1.20 | 1.03 | 0.30 | 0.61 | 0.42 | 0.77 | 0.73 |
| Dataset 1 | Dataset 5 | 2.79 | 3.35 | 0.30 | 0.45 | 0.84 | 0.99 | 1.79 |
| Dataset 2 | Dataset 1 | 0.97 | 0.97 | 0.31 | 0.30 | 0.28 | 0.17 | 0.13 |
| Dataset 2 | Dataset 3 | 0.71 | 0.68 | 0.31 | 0.17 | 0.23 | 0.10 | 0.12 |
| Dataset 2 | Dataset 4, Day -1 | 0.98 | 1.14 | 0.31 | 0.61 | 0.57 | 0.97 | 0.85 |
| Dataset 2 | Dataset 4, Day -3 | 1.35 | 1.10 | 0.31 | 0.61 | 0.39 | 0.62 | 0.39 |
| Dataset 2 | Dataset 5 | 1.75 | 1.51 | 0.31 | 0.45 | 0.21 | 0.01 | 0.04 |
| Dataset 3 | Dataset 1 | 1.06 | 0.97 | 0.17 | 0.30 | 0.18 | 0.30 | 0.03 |
| Dataset 3 | Dataset 2 | 0.84 | 0.79 | 0.17 | 0.31 | 0.23 | 0.41 | 0.43 |
| Dataset 3 | Dataset 4, Day -1 | 1.25 | 1.18 | 0.17 | 0.61 | 0.39 | 0.99 | 1.09 |
| Dataset 3 | Dataset 4, Day -3 | 1.77 | 1.12 | 0.17 | 0.61 | 0.26 | 0.77 | 0.18 |
| Dataset 3 | Dataset 5 | 1.92 | 1.30 | 0.17 | 0.45 | 0.15 | 0.06 | -0.05 |
| Dataset 4, Day -3 | Dataset 1 | 1.05 | 0.96 | 0.61 | 0.30 | 0.42 | 0.14 | 0.11 |
| Dataset 4, Day -3 | Dataset 2 (Age < 16) | 1.07 | 0.98 | 0.61 | 0.37 | 0.47 | 0.35 | 0.27 |
| Dataset 4, Day -3 | Dataset 3 (Age < 16) | 1.20 | 0.74 | 0.61 | 0.14 | 0.37 | 0.12 | 0.02 |
| Dataset 4, Day -3 | Dataset 5 | 5.33 | 4.72 | 0.61 | 0.45 | 0.95 | 0.99 | 1.20 |
| Dataset 4, Day -1 | Dataset 1 | 0.99 | 1.26 | 0.61 | 0.30 | 0.26 | 0.01 | -0.15 |
| Dataset 4, Day -1 | Dataset 2 (Age < 16) | 1.02 | 1.07 | 0.61 | 0.37 | 0.38 | 0.23 | 0.14 |
| Dataset 4, Day -1 | Dataset 3 (Age < 16) | 0.88 | 0.87 | 0.61 | 0.14 | 0.22 | 0.01 | -0.20 |
| Dataset 4, Day -1 | Dataset 5 | 2.63 | 2.29 | 0.61 | 0.45 | 0.75 | 0.85 | 0.83 |
| Dataset 5 | Dataset 1 | 1.04 | 0.98 | 0.45 | 0.30 | 0.40 | 0.10 | 0.26 |
| Dataset 5 | Dataset 2 | 1.05 | 0.91 | 0.45 | 0.31 | 0.49 | 0.57 | 0.77 |
| Dataset 5 | Dataset 3 | 1.06 | 0.69 | 0.45 | 0.17 | 0.41 | 0.29 | 0.42 |
| Dataset 5 | Dataset 4, Day -1 | 1.05 | 1.12 | 0.45 | 0.61 | 0.60 | 0.99 | 1.28 |
| Dataset 5 | Dataset 4, Day -3 | 1.20 | 1.14 | 0.45 | 0.61 | 0.49 | 0.69 | 0.78 |

**Supplementary Table 4.** In-sample and generalizability performance metrics for logistic regression with each training/test pair, Alternative Subset 1. Model denotes whether the model evaluated is the original model (age, WBC, and PLT), the full model (using all available variables in the subset), or the model chosen by best subsets selection (BSS) with that training data. Missing values in the PPV indicate that there were 0 predicted positive cases.

| Training data | Test data | Model | Sensitivity | Specificity | PPV | NPV | AUC |
| --- | --- | --- | --- | --- | --- | --- | --- |
| Dataset 1 | Dataset 1 | Original | 0.742 (0.721, 0.763) | 0.757 (0.743, 0.77) | 0.563 (0.542, 0.583) | 0.874 (0.863, 0.885) | 0.83 (0.81, 0.84) |
| Dataset 1 | Dataset 1 | Full | 0.767 (0.747, 0.787) | 0.775 (0.762, 0.788) | 0.59 (0.57, 0.611) | 0.888 (0.877, 0.898) | 0.85 (0.84, 0.86) |
| Dataset 1 | Dataset 1 | BSS | 0.767 (0.747, 0.787) | 0.778 (0.765, 0.791) | 0.593 (0.572, 0.614) | 0.888 (0.877, 0.898) | 0.85 (0.84, 0.86) |
| Dataset 1 | Dataset 3 | Original | 0.914 (0.873, 0.945) | 0.367 (0.341, 0.395) | 0.228 (0.203, 0.255) | 0.954 (0.932, 0.971) | 0.669 (0.639, 0.698) |
| Dataset 1 | Dataset 3 | Full | 0 (0, 0.014) | 0.999 (0.996, 1) | 0 (0, 0.975) | 0.83 (0.81, 0.849) | 0.679 (0.652, 0.706) |
| Dataset 1 | Dataset 3 | BSS | 0 (0, 0.014) | 0.999 (0.996, 1) | 0 (0, 0.975) | 0.83 (0.81, 0.849) | 0.693 (0.665, 0.722) |
| Dataset 1 | Dataset 3 (Age < 16) | Original | 0.759 (0.653, 0.846) | 0.751 (0.709, 0.789) | 0.348 (0.279, 0.422) | 0.947 (0.919, 0.967) | 0.83 (0.782, 0.879) |
| Dataset 1 | Dataset 3 (Age < 16) | Full | 0 (0, 0.043) | 1 (0.992, 1) |  | 0.851 (0.818, 0.879) | 0.434 (0.381, 0.487) |
| Dataset 1 | Dataset 3 (Age < 16) | BSS | 0 (0, 0.043) | 1 (0.992, 1) |  | 0.851 (0.818, 0.879) | 0.425 (0.37, 0.48) |
| Dataset 1 | Dataset 5 | Original | 0.994 (0.967, 1) | 0.062 (0.032, 0.105) | 0.474 (0.421, 0.528) | 0.923 (0.64, 0.998) | 0.67 (0.614, 0.726) |
| Dataset 1 | Dataset 5 | Full | 0 (0, 0.022) | 1 (0.981, 1) |  | 0.54 (0.487, 0.592) | 0.474 (0.428, 0.52) |
| Dataset 1 | Dataset 5 | BSS | 0 (0, 0.022) | 1 (0.981, 1) |  | 0.54 (0.487, 0.592) | 0.463 (0.413, 0.514) |
| Dataset 3 | Dataset 3 | Original | 0.789 (0.734, 0.837) | 0.776 (0.751, 0.798) | 0.418 (0.374, 0.464) | 0.947 (0.932, 0.96) | 0.85 (0.82, 0.87) |
| Dataset 3 | Dataset 3 | Full | 0.773 (0.717, 0.823) | 0.801 (0.778, 0.823) | 0.443 (0.396, 0.49) | 0.945 (0.93, 0.958) | 0.87 (0.84, 0.89) |
| Dataset 3 | Dataset 3 | BSS | 0.773 (0.717, 0.823) | 0.8 (0.776, 0.821) | 0.441 (0.394, 0.488) | 0.945 (0.93, 0.958) | 0.87 (0.84, 0.89) |
| Dataset 3 | Dataset 1 | Original | 0.695 (0.672, 0.717) | 0.756 (0.743, 0.769) | 0.546 (0.525, 0.567) | 0.855 (0.843, 0.866) | 0.803 (0.79, 0.815) |
| Dataset 3 | Dataset 1 | Full | 0.959 (0.949, 0.968) | 0.253 (0.239, 0.266) | 0.351 (0.337, 0.365) | 0.936 (0.92, 0.95) | 0.805 (0.793, 0.818) |
| Dataset 3 | Dataset 1 | BSS | 0.942 (0.929, 0.952) | 0.328 (0.314, 0.343) | 0.372 (0.357, 0.386) | 0.93 (0.916, 0.943) | 0.813 (0.8, 0.825) |
| Dataset 3 | Dataset 5 | Original | 0.44 (0.363, 0.519) | 0.882 (0.828, 0.924) | 0.76 (0.663, 0.842) | 0.649 (0.588, 0.706) | 0.74 (0.688, 0.792) |
| Dataset 3 | Dataset 5 | Full | 0.596 (0.518, 0.672) | 0.749 (0.682, 0.808) | 0.669 (0.587, 0.744) | 0.685 (0.618, 0.747) | 0.715 (0.661, 0.768) |
| Dataset 3 | Dataset 5 | BSS | 0.578 (0.499, 0.654) | 0.779 (0.715, 0.836) | 0.691 (0.607, 0.766) | 0.685 (0.619, 0.745) | 0.724 (0.67, 0.777) |
| Dataset 5 | Dataset 5 | Original | 0.693 (0.617, 0.762) | 0.641 (0.569, 0.708) | 0.622 (0.548, 0.692) | 0.71 (0.637, 0.776) | 0.73 (0.68, 0.79) |
| Dataset 5 | Dataset 5 | Full | 0.711 (0.636, 0.778) | 0.641 (0.569, 0.708) | 0.628 (0.554, 0.697) | 0.723 (0.649, 0.788) | 0.72 (0.66, 0.77) |
| Dataset 5 | Dataset 5 | BSS | 0.717 (0.642, 0.784) | 0.636 (0.564, 0.703) | 0.626 (0.553, 0.695) | 0.725 (0.652, 0.791) | 0.74 (0.69, 0.79) |
| Dataset 5 | Dataset 1 | Original | 0.777 (0.756, 0.796) | 0.68 (0.665, 0.694) | 0.506 (0.486, 0.525) | 0.878 (0.866, 0.89) | 0.809 (0.797, 0.821) |
| Dataset 5 | Dataset 1 | Full | 0.852 (0.834, 0.868) | 0.526 (0.511, 0.542) | 0.431 (0.414, 0.448) | 0.894 (0.881, 0.906) | 0.8 (0.788, 0.813) |
| Dataset 5 | Dataset 1 | BSS | 0.753 (0.732, 0.774) | 0.675 (0.66, 0.689) | 0.494 (0.475, 0.514) | 0.866 (0.854, 0.878) | 0.786 (0.774, 0.799) |
| Dataset 5 | Dataset 3 | Original | 0.863 (0.815, 0.903) | 0.593 (0.565, 0.62) | 0.302 (0.269, 0.337) | 0.955 (0.938, 0.968) | 0.836 (0.81, 0.863) |
| Dataset 5 | Dataset 3 | Full | 0.859 (0.811, 0.9) | 0.597 (0.57, 0.625) | 0.304 (0.271, 0.339) | 0.954 (0.937, 0.968) | 0.824 (0.796, 0.853) |
| Dataset 5 | Dataset 3 | BSS | 0.875 (0.828, 0.913) | 0.598 (0.57, 0.626) | 0.308 (0.275, 0.343) | 0.959 (0.943, 0.972) | 0.837 (0.81, 0.864) |
| Dataset 5 | Dataset 3 (Age > 16) | Original | 0.939 (0.89, 0.97) | 0.522 (0.486, 0.558) | 0.298 (0.258, 0.339) | 0.975 (0.955, 0.988) | 0.861 (0.832, 0.89) |
| Dataset 5 | Dataset 3 (Age > 16) | Full | 0.957 (0.914, 0.983) | 0.527 (0.491, 0.563) | 0.304 (0.265, 0.346) | 0.983 (0.965, 0.993) | 0.867 (0.839, 0.896) |
| Dataset 5 | Dataset 3 (Age > 16) | BSS | 0.945 (0.898, 0.974) | 0.574 (0.537, 0.609) | 0.324 (0.282, 0.368) | 0.98 (0.962, 0.991) | 0.884 (0.857, 0.912) |

**Supplementary Table 5.** In-sample and generalizability performance metrics for logistic regression with each training/test pair, Alternative Subset 2. Model denotes whether the model evaluated is the original model (age, WBC, and PLT), the full model (using all available variables in the subset), or the model chosen by best subsets selection (BSS) with that training data.

| Training data | Test data | Model | Sensitivity | Specificity | PPV | NPV | AUC |
| --- | --- | --- | --- | --- | --- | --- | --- |
| Dataset 2 | Dataset 2 | Original | 0.805 (0.762, 0.844) | 0.816 (0.786, 0.844) | 0.693 (0.647, 0.736) | 0.891 (0.865, 0.913) | 0.88 (0.86, 0.9) |
| Dataset 2 | Dataset 2 | Full | 0.789 (0.745, 0.83) | 0.818 (0.788, 0.845) | 0.69 (0.644, 0.733) | 0.883 (0.856, 0.906) | 0.9 (0.88, 0.91) |
| Dataset 2 | Dataset 2 | BSS | 0.803 (0.759, 0.842) | 0.822 (0.792, 0.849) | 0.698 (0.653, 0.741) | 0.89 (0.864, 0.913) | 0.9 (0.88, 0.92) |
| Dataset 2 | Dataset 3 | Original | 0.633 (0.571, 0.692) | 0.853 (0.833, 0.872) | 0.46 (0.407, 0.514) | 0.922 (0.905, 0.936) | 0.852 (0.828, 0.877) |
| Dataset 2 | Dataset 3 | Full | 0.695 (0.635, 0.751) | 0.859 (0.838, 0.877) | 0.493 (0.44, 0.546) | 0.934 (0.919, 0.948) | 0.863 (0.839, 0.887) |
| Dataset 2 | Dataset 3 | BSS | 0.711 (0.651, 0.766) | 0.858 (0.838, 0.876) | 0.497 (0.445, 0.55) | 0.938 (0.922, 0.951) | 0.862 (0.838, 0.886) |
| Dataset 2 | Dataset 5 | Original | 0.341 (0.27, 0.419) | 0.949 (0.909, 0.975) | 0.851 (0.743, 0.926) | 0.63 (0.572, 0.685) | 0.734 (0.682, 0.786) |
| Dataset 2 | Dataset 5 | Full | 0.467 (0.39, 0.546) | 0.868 (0.813, 0.912) | 0.75 (0.656, 0.83) | 0.658 (0.597, 0.715) | 0.716 (0.663, 0.769) |
| Dataset 2 | Dataset 5 | BSS | 0.467 (0.39, 0.546) | 0.863 (0.807, 0.908) | 0.743 (0.648, 0.823) | 0.656 (0.595, 0.714) | 0.718 (0.665, 0.771) |
| Dataset 3 | Dataset 3 | Original | 0.777 (0.721, 0.827) | 0.779 (0.755, 0.801) | 0.41 (0.366, 0.456) | 0.946 (0.931, 0.959) | 0.85 (0.83, 0.87) |
| Dataset 3 | Dataset 3 | Full | 0.77 (0.713, 0.82) | 0.807 (0.784, 0.828) | 0.441 (0.394, 0.488) | 0.947 (0.932, 0.959) | 0.86 (0.84, 0.89) |
| Dataset 3 | Dataset 3 | BSS | 0.77 (0.713, 0.82) | 0.81 (0.787, 0.831) | 0.445 (0.398, 0.492) | 0.947 (0.932, 0.959) | 0.86 (0.84, 0.89) |
| Dataset 3 | Dataset 2 | Original | 0.891 (0.855, 0.92) | 0.723 (0.689, 0.755) | 0.623 (0.581, 0.664) | 0.928 (0.903, 0.948) | 0.892 (0.872, 0.912) |
| Dataset 3 | Dataset 2 | Full | 0.824 (0.782, 0.861) | 0.813 (0.783, 0.841) | 0.694 (0.649, 0.737) | 0.9 (0.874, 0.922) | 0.9 (0.881, 0.919) |
| Dataset 3 | Dataset 2 | BSS | 0.832 (0.79, 0.868) | 0.822 (0.792, 0.849) | 0.706 (0.661, 0.748) | 0.905 (0.88, 0.926) | 0.9 (0.88, 0.919) |
| Dataset 3 | Dataset 5 | Original | 0.425 (0.349, 0.504) | 0.883 (0.83, 0.925) | 0.755 (0.656, 0.838) | 0.644 (0.584, 0.702) | 0.741 (0.69, 0.793) |
| Dataset 3 | Dataset 5 | Full | 0.551 (0.472, 0.628) | 0.787 (0.723, 0.842) | 0.687 (0.601, 0.764) | 0.674 (0.609, 0.734) | 0.719 (0.666, 0.772) |
| Dataset 3 | Dataset 5 | BSS | 0.545 (0.466, 0.622) | 0.782 (0.717, 0.837) | 0.679 (0.593, 0.757) | 0.67 (0.605, 0.73) | 0.722 (0.669, 0.774) |
| Dataset 5 | Dataset 5 | Original | 0.713 (0.638, 0.78) | 0.609 (0.537, 0.678) | 0.607 (0.535, 0.676) | 0.714 (0.64, 0.781) | 0.74 (0.69, 0.79) |
| Dataset 5 | Dataset 5 | Full | 0.683 (0.606, 0.752) | 0.64 (0.568, 0.707) | 0.616 (0.542, 0.687) | 0.704 (0.631, 0.77) | 0.72 (0.67, 0.77) |
| Dataset 5 | Dataset 5 | BSS | 0.701 (0.625, 0.769) | 0.624 (0.553, 0.692) | 0.613 (0.54, 0.682) | 0.711 (0.637, 0.777) | 0.74 (0.68, 0.79) |
| Dataset 5 | Dataset 2 | Original | 0.952 (0.925, 0.971) | 0.514 (0.477, 0.551) | 0.502 (0.465, 0.539) | 0.954 (0.929, 0.973) | 0.869 (0.847, 0.891) |
| Dataset 5 | Dataset 2 | Full | 0.965 (0.941, 0.981) | 0.473 (0.436, 0.51) | 0.485 (0.449, 0.522) | 0.964 (0.939, 0.981) | 0.851 (0.828, 0.874) |
| Dataset 5 | Dataset 2 | BSS | 0.96 (0.935, 0.977) | 0.469 (0.432, 0.506) | 0.482 (0.446, 0.518) | 0.958 (0.932, 0.976) | 0.858 (0.835, 0.88) |
| Dataset 5 | Dataset 2 (Age > 16) | Original | 0.962 (0.93, 0.983) | 0.496 (0.455, 0.538) | 0.446 (0.402, 0.49) | 0.969 (0.942, 0.986) | 0.882 (0.858, 0.907) |
| Dataset 5 | Dataset 2 (Age > 16) | Full | 0.979 (0.952, 0.993) | 0.458 (0.416, 0.5) | 0.432 (0.39, 0.475) | 0.981 (0.957, 0.994) | 0.858 (0.831, 0.884) |
| Dataset 5 | Dataset 2 (Age > 16) | BSS | 0.967 (0.935, 0.985) | 0.467 (0.425, 0.509) | 0.433 (0.39, 0.476) | 0.971 (0.943, 0.987) | 0.864 (0.839, 0.89) |
| Dataset 5 | Dataset 3 | Original | 0.871 (0.824, 0.91) | 0.591 (0.564, 0.618) | 0.297 (0.264, 0.331) | 0.959 (0.942, 0.971) | 0.838 (0.811, 0.865) |
| Dataset 5 | Dataset 3 | Full | 0.875 (0.828, 0.913) | 0.582 (0.554, 0.609) | 0.293 (0.261, 0.326) | 0.959 (0.943, 0.972) | 0.838 (0.811, 0.865) |
| Dataset 5 | Dataset 3 | BSS | 0.883 (0.837, 0.92) | 0.566 (0.538, 0.593) | 0.287 (0.255, 0.32) | 0.961 (0.944, 0.973) | 0.842 (0.815, 0.868) |
| Dataset 5 | Dataset 3 (Age > 16) | Original | 0.939 (0.89, 0.97) | 0.525 (0.489, 0.56) | 0.292 (0.253, 0.333) | 0.976 (0.957, 0.989) | 0.864 (0.835, 0.892) |
| Dataset 5 | Dataset 3 (Age > 16) | Full | 0.945 (0.898, 0.974) | 0.529 (0.493, 0.564) | 0.295 (0.256, 0.336) | 0.979 (0.96, 0.99) | 0.88 (0.852, 0.907) |
| Dataset 5 | Dataset 3 (Age > 16) | BSS | 0.945 (0.898, 0.974) | 0.528 (0.492, 0.563) | 0.294 (0.256, 0.336) | 0.979 (0.96, 0.99) | 0.886 (0.859, 0.913) |

**Supplementary Table 6.**
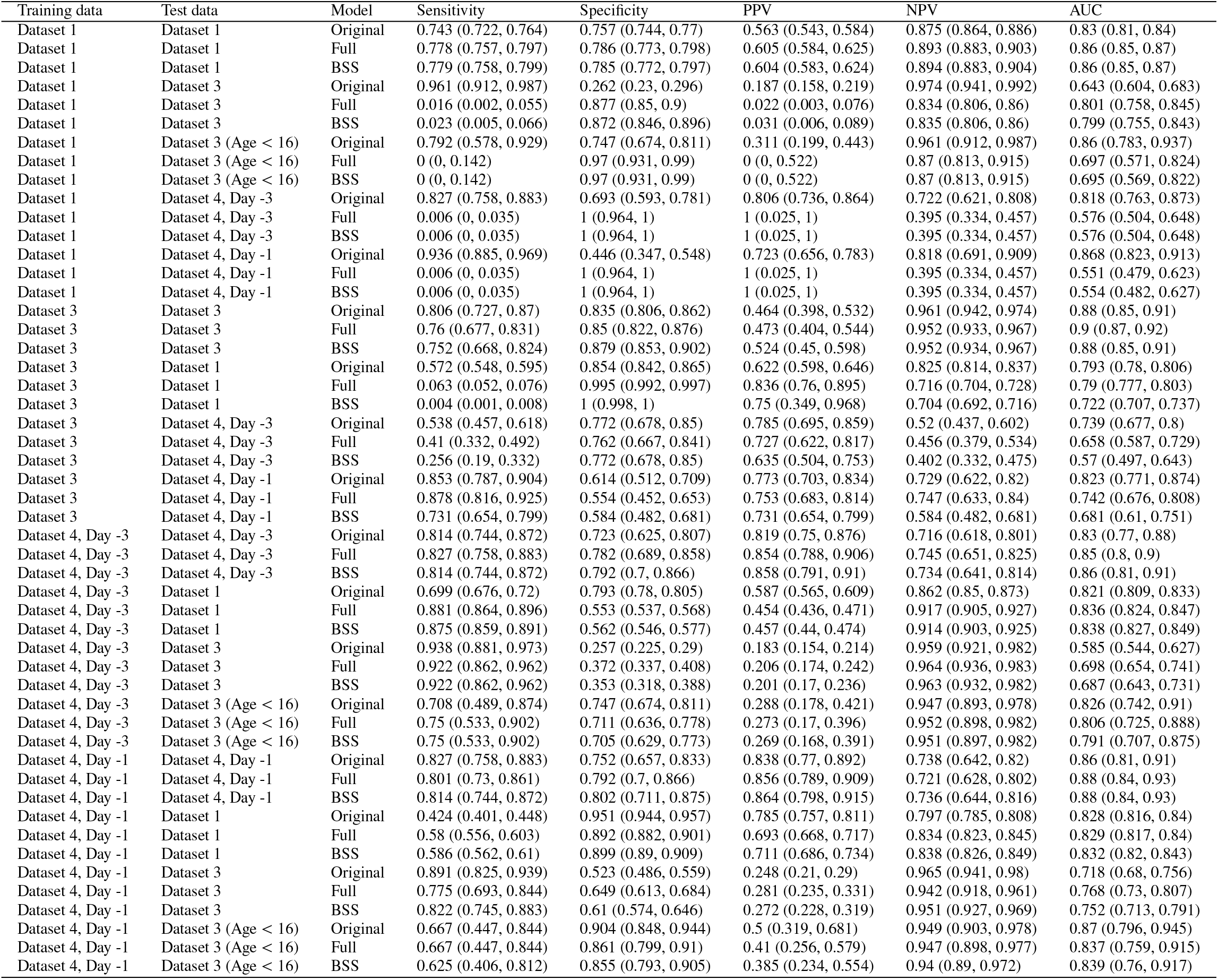
In-sample and generalizability performance metrics for logistic regression with each training/test pair, Alternative Subset 3. Model denotes whether the model evaluated is the original model (age, WBC, and PLT), the full model (using all available variables in the subset), or the model chosen by best subsets selection (BSS) with that training data.

